# The Structural Basis of Malodorant Skatole Formation by the Glycyl Radical Enzyme Indoleacetate Decarboxylase

**DOI:** 10.64898/2026.05.24.727522

**Authors:** Christa N. Imrich, Lindsey R.F. Backman, Abigail P. Allworth, Mary C. Andorfer, Jared C. Paris, Nina M. Greeley, Beverly Fu, Emily P. Balskus, Catherine L. Drennan

## Abstract

Glycyl radical enzymes (GREs) catalyze challenging chemical reactions using a post-translationally installed glycyl radical cofactor. One such enzyme, indoleacetate decarboxylase (IAD), performs the radical-based decarboxylation of indole-3-acetate (I3A) to form the malodorant molecule skatole. In addition to being an odor nuisance, skatole is a human and livestock lung toxin, a suspected carcinogen, and a mosquito attractant, all of which impact human health, agriculture, food production, and wastewater treatment. Here, we use cryogenic electron microscopy to solve a 2.45-Å resolution structure of indoleacetate decarboxylase from the gut bacterium *Olsenella uli*. We observe IAD in a homotetrameric form with the substrate I3A bound in all four protomers. The positioning of I3A in the active site is unexpected and is more consistent with a Kolbe-type decarboxylation mechanism, i.e. a decarboxylation initiated by a 1-electron oxidation of the carboxylate moiety rather than being initiated by hydrogen atom transfer (HAT). Previously, a high deuterium content in skatole from IAD assays in D_2_O was used to support a HAT mechanism over a Kolbe-type mechanism. However, we show here that deuterium content does not necessarily inform on mechanism as IAD can catalyze the exchange of skatole’s 3′-methyl hydrogens post-turnover. Structural comparisons show that both IAD and HPAD display structural features that are not found in other characterized GREs, suggesting that they represent a distinct GRE-subclass. Collectively, these insights will inform IAD inhibitor design aimed at decreasing skatole production.

**Significance Statement:** Glycyl radical enzymes (GREs) are a superfamily of enzymes that catalyze challenging chemical reactions in anaerobic environments. One such enzyme, indoleacetate decarboxylase (IAD) performs a C–C bond cleavage and decarboxylation reaction on the tryptophan metabolite indole-3-acetate (I3A). Decarboxylation of I3A by IAD forms the malodorant molecule skatole which negatively impacts human health, livestock health, odor emissions, the environment, and food production. Here, we present the first structure of IAD with I3A bound, revealing the substrate binding mode and active site architecture. This work provides new insight into the catalytic mechanism of the enzyme and illuminates new avenues for control of the odor nuisance skatole.

## Introduction

3-methylindole, commonly known as skatole, is a malodorous compound produced by microbes (1). It is primarily responsible for the distinct odor of mammalian excrement. Humans have a low threshold of detection for skatole and can identify its odor at nanomolar concentrations in water and picomolar concentrations in the air (2, 3). Consequently, skatole impacts zoning laws for wastewater treatment sites and agricultural farms (4). Mosquitos, like humans, possess odorant receptors for skatole and recent work demonstrated that *Culex* mosquitos are narrowly tuned to skatole, which functions as an oviposition attractant, directing female mosquitos to egg-laying sites (5). Skatole is produced in the GI tract of livestock such as pigs, and in male pigs, it can accumulate in fatty tissues, contributing to a phenomenon known as boar taint (6, 7). Boar taint causes malodor in pork meat that then must be thrown out despite being of otherwise adequate quality for consumption, resulting in significant food waste (8). Further, skatole can be converted to 3-methyleneindolenine in the lungs of both humans and livestock by cytochrome P450s (9). 3-methyleneindolenine is a toxin suspected to interfere with cell adhesion in the lung epithelium, and causes a disease known as fog fever. In livestock the condition can be severe, and up to 50% of cattle afflicted with the disease will die (10). Skatole has demonstrably far-ranging implications in urbanism, human health and disease, agriculture, and the environment. It is therefore desirable to understand the molecular basis for production of skatole such that remediation efforts for its impacts may be designed, tested, and employed.

Until recently, the pathway or enzymes involved in the microbial synthesis of skatole were unknown. However, in 2018, Liu *et al.* reported that a glycyl radical enzyme (GRE) forms skatole via a decarboxylation reaction (11). GREs are strictly anaerobic enzymes that utilize an oxygen-sensitive glycyl radical cofactor to perform catalysis (12). This radical cofactor is post-translationally installed by a cognate glycyl radical enzyme activating enzyme (GRE-AE) that is specific to each GRE. GRE-AEs are members of the radical *S*-adenosyl-L-methionine (RS) superfamily of enzymes (13). Therefore, GRE-AEs contain a canonical RS domain that houses a [4Fe–4S]^2+/+^ cluster coordinated by a CX_3_CX_2_C motif and a molecule of *S*-adenosyl-L-methionine (AdoMet). Upon reduction of the iron-sulfur cluster by a biological electron donor, GRE-AEs reductively cleave AdoMet to form the highly reactive 5’dAdo• radical species and methionine (14) (**Fig 1A**). This 5’dAdo• radical abstracts a hydrogen atom from a conserved glycine residue at the C-terminus of the GRE polypeptide found in the fingerprint motif RVx**G**[FWY]x_6-8_[FL]x_4_Qx_2_[IV]x_2_R (15), thereby producing a stable glycyl radical that can be used for many rounds of catalytic turnover (**Fig 1A**). The catalytic cycle of GREs share certain conserved features (12). To initiate catalysis, the glycyl radical abstracts a hydrogen atom from the thiol of a conserved cysteine residue in the GRE polypeptide, forming a transient thiyl radical species. This highly oxidative thiyl radical then acts on substrate, typically by abstracting a hydrogen atom from substrate to form a substrate radical. The substrate radical can rearrange to form a product radical, and the final step is the re-abstraction of the cysteine thiol hydrogen atom by the product radical, reforming the thiyl radical and forming product (**Fig 1A**). The thiyl radical can then reform glycyl radical and the cycle can begin again for subsequent rounds of turnover.

**Figure 1.**
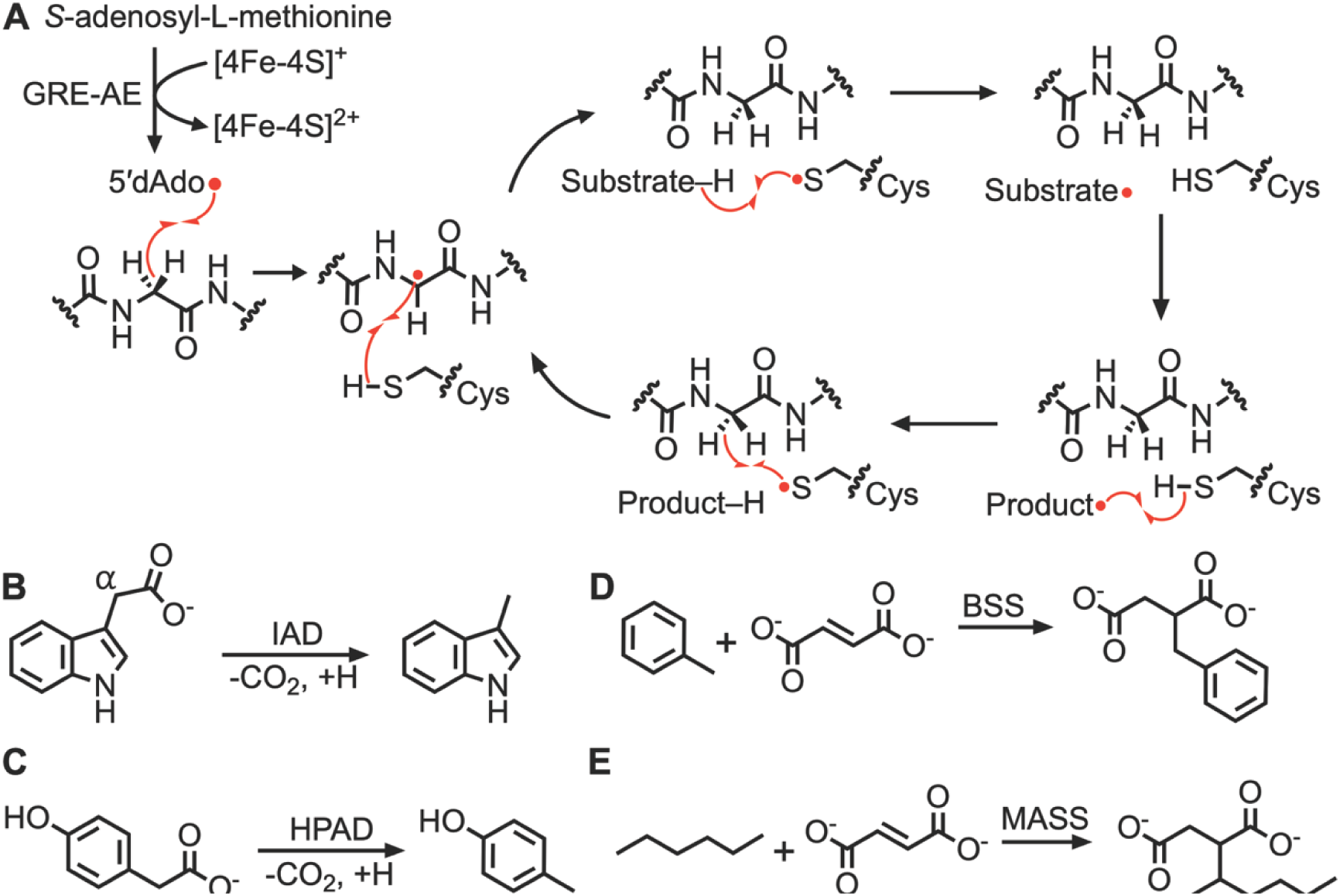
Post-translational radical installation on the GRE and the conserved mechanism of GRE catalysis. **A.** AdoMet is reductively cleaved by the GRE-AE using a [4Fe–4S]^+^ cluster to form the highly reactive 5’dAdo radical. The 5’dAdo radical abstracts a hydrogen atom from the conserved glycine to form the glycyl radical. Once the glycyl radical is formed, the cofactor can be used for many rounds of catalysis via a transient thiyl radical formed on the conserved cysteine. **B.** The reaction of indoleacetate decarboxylase. **C.** The reaction of 4-hydroxyphenylacetate decarboxylase **D.** The reaction of benzylsuccinate synthase. **E.** The reaction of 1-methylalkylsuccinate synthase.

The skatole-forming GRE was aptly named indoleacetate decarboxylase (IAD). It catalyzes the radical-dependent decarboxylation of indole-3-acetate (I3A) (**Fig 1B**) (11). Thus, IAD is codified as a member of a subclass of GREs known as the GRE decarboxylases. The most well-characterized and founding member of the GRE decarboxylases, hydroxyphenylacetate decarboxylase (HPAD), catalyzes a similar decarboxylation of 4-hydroxyphenylacetate (4HP) to form *p*-cresol, a bacteriostatic molecule (16) (**Fig 1C**). At the time of IAD’s discovery, HPAD was the only GRE decarboxylase with an experimentally solved structure and remains the only high-quality GRE decarboxylase structure to date (17). X-ray crystal structures of HPAD revealed a heterooctamer (a homotetramer of βγ heterodimers), which added HPAD to a subset of GREs that are multi-subunit. Benzylsuccinate synthase (BSS) and 1-methylalkylsuccinate synthase (MASS), both members of the X-succinate synthase subclass of GREs, represent the other known multi-subunit GREs (18–20). BSS catalyzes the C–H bond activation of toluene, adding it to fumarate to form benzylsuccinate (**Fig 1D**), whereas MASS catalyzes C–H bond activation of *n*-alkanes, adding them to fumarate to form alkylsuccinates (**Fig 1E**). Despite architectural variation across GREs such as the existence of small subunits, most GREs employ a hydrogen atom transfer (HAT) from substrate as the initial step in the catalytic cycle. However, HPAD is the sole GRE proposed to use an alternative mechanism–a Kolbe-type decarboxylation–that begins with an electron transfer from the substrate carboxylate to the thiyl radical. Initial biochemical studies of IAD seemed to support a canonical GRE mechanism that begins with HAT (21), in conflict with the mechanism proposed for HPAD. It is not clear why two enzymes that perform remarkably similar chemical reactions would use distinctly different mechanisms, and we wanted to further interrogate whether the mechanisms are the same or different, and if so, if the IAD structure can explain why. Here we present a high-resolution cryogenic electron microscopy (cryo-EM) structure of IAD in a substrate-bound state and perform biochemical studies to further probe the enigmatic mechanism of IAD catalysis. Our findings suggest that IAD and HPAD both likely use the same mechanism, a Kolbe-type decarboxylation.

## Results

### The SPT Labtech chameleon enabled the capture of a high-resolution structure of IAD in a tetrameric state

We determined the structure of IAD from *Olsenella uli* DSM 7084 (*Ou*IAD) to 2.45-Å resolution by cryo-EM (**Fig S1, S2, Table S1**). Key to the obtainment of this high-resolution structure was the use of a SPT Labtech chameleon^®^ for grid preparation. Consistent with previous observations, grid preparation using the chameleon^®^ provided more intact particles and ameliorated air-water interface denaturation (22, 23). Surprisingly, we observed IAD in a homotetrameric state (**Fig 2A**). The canonical GRE dimer is observed within our structure (**Fig 2B),** and two of these dimers come together to form the homotetramer. Further, this IAD tetramer is consistent with GRE tetramers that have been observed in the crystal lattices of GRE structures solved by X-ray crystallography (**Fig S3**). Each IAD monomer maintains a common GRE fold consisting of a 10-stranded β-barrel surrounded by α helices (**Fig 2C**). The four protomers are highly similar, with an r.m.s.d. of 0.17 Å on average between Cα atoms of the polypeptides. Two β hairpin loops, known as the Gly and Cys loops, meet in the center of the barrel and compose part of the buried active site. The Gly loop is part of a flexible C-terminal domain called the glycyl radical domain (GRD) that houses the catalytically essential glycine residue (Gly853 in *Ou*IAD) (**Fig 2C**). Likewise, the Cys loop contains a catalytically essential cysteine residue (Cys 500 in *Ou*IAD). In our structure, Gly853 and Cys500 are positioned appropriately to generate a transient thiyl radical for catalysis, with a Gly-Cα to Cys-Sγ distance of 4.6 Å (**Fig 2C, inset**).

**Figure 2.**
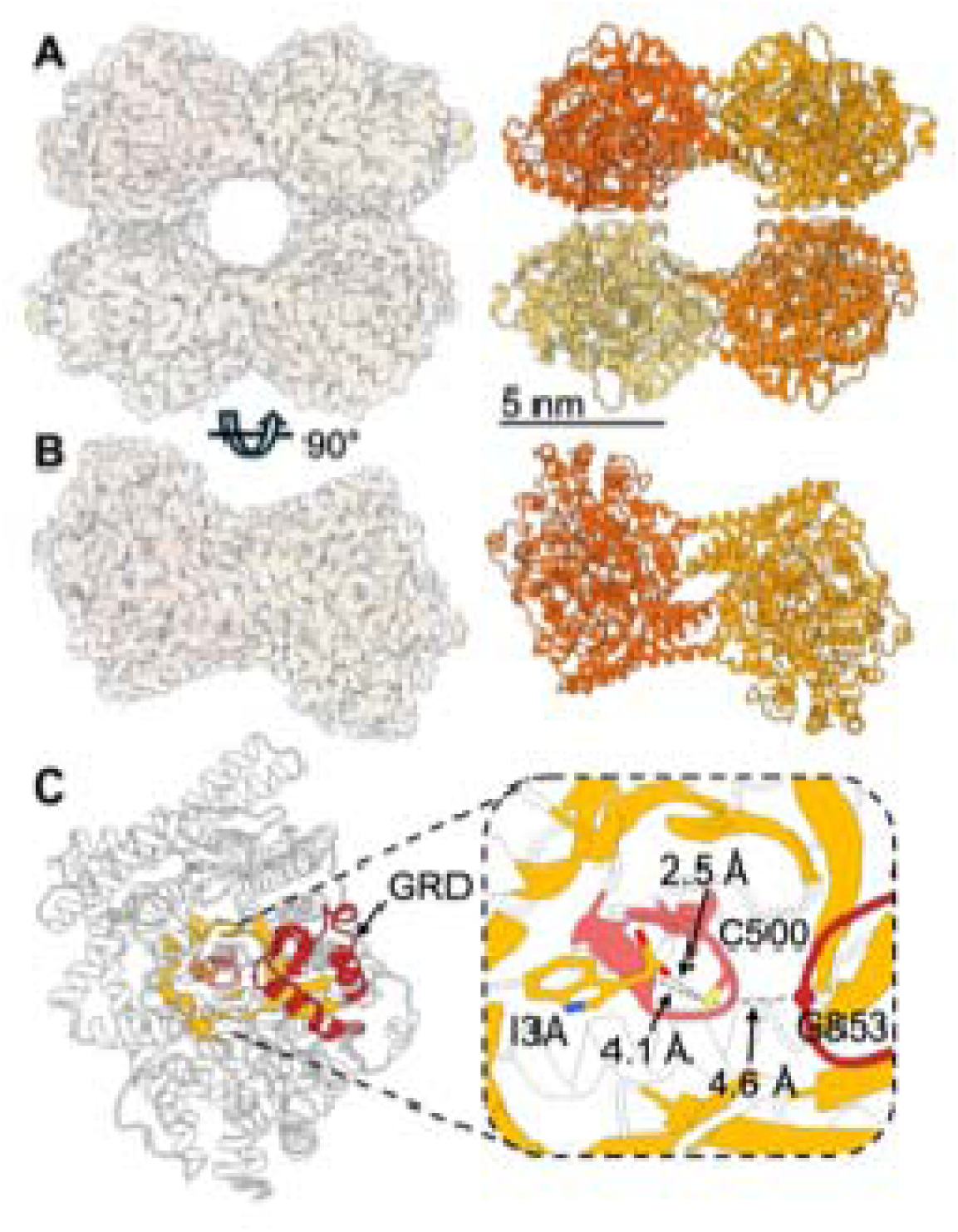
A 2.45-Å resolution structure of *Ou*IAD shows a conserved GRE architecture. **A.** The 3D reconstruction to 2.45-Å resolution and corresponding *Ou*IAD model shows a tetrameric species. **B.** A canonical GRE dimer is maintained within the *Ou*IAD tetramer. **C.** *Ou*IAD maintains a conserved GRE fold overall consisting of two 5-stranded β-sheets that are antiparallel to one another and come together to form a complete 10-stranded β-barrel surrounded by α helices. GRD shown in firebrick. Inset: the Gly and Cys loops meet in the middle of the barrel and are positioned appropriately for radical catalysis to proceed.

### An IAD tetramer can be observed by mass photometry

IAD has been observed by size exclusion chromatography in mixed oligomeric states, with dimer and monomer populations, in addition to some higher-order oligomeric states (11, 21) (**Fig S4**). Therefore, we set out to use mass photometry to probe the oligomeric state distribution of IAD in solution. By mass photometry analysis, we observed a concentration-dependent shift toward higher oligomeric state species regardless of the presence of IAD’s I3A substrate (**Fig S5A-B**). At IAD concentrations as low as 100-200 nM, we observe an appreciable peak shoulder near 400 kDa, the approximate molecular weight of an IAD tetramer, in addition to the diminishing of a peak near 100 kDa (**Fig S5A-B**), the approximate molecular weight of an IAD monomer. The concentration of IAD at which we begin to observe tetramer by mass photometry (100 nM) is 1000 times more dilute than the concentration of IAD (100 µM) used to prepare the cryo-EM grid that resulted in the tetrameric structure. Collectively, these data strongly indicate that the tetrameric state in addition to the dimeric state are relevant oligomeric states for this GRE.

### IAD shares highest structural similarity with small subunit-containing GREs

IAD shares the highest structural homology with the three structurally characterized GREs that employ small subunits: HPAD, BSS, and MASS (r.m.s.d. of 1.08 Å, 1.15 Å, 1.14 Å between core Cα atoms). Although IAD does not have small subunits, it does have multiple extended regions, which include an N-terminal extension (residues 4-41), an extended region near the active site (residues 124-169, aka extension 1), and an elongated loop (residues 651-675, aka extension 2) (**Fig 3A, S6**). N-terminal extensions are also found in BSS and MASS, but they are shorter than IAD’s N-terminus, which covers ∼1750 Å^2^ of the edge surface of each protomer (**Fig 3A-D**). In HPAD, the N-terminal 28 residues are disordered, so the degree of similarity to the N-terminus to IAD is not known (**Fig 3B**). IAD’s extension 1 is located N-terminal to the 10-stranded barrel, preceding the conserved helices that precede β1 (**Fig S6**), and is adjacent to the so-called cap helix (residues 390-396 in IAD) that sits over GRE active sites. The second extension is found between a set of two conserved helices that lie in between strands β7 and β8 (**Fig S6**). HPAD shares extensions 1 and 2 with IAD (**Fig 3A-B, Fig S7**). Sequence alignment of IAD, HPAD, and the other known GRE decarboxylases, phenylacetate decarboxylase (PAD) and arylacetate decarboxylase (AAD), shows that AAD and PAD do not appear to have the extensions observed in IAD and HPAD (**Fig S7**).

**Figure 3.**
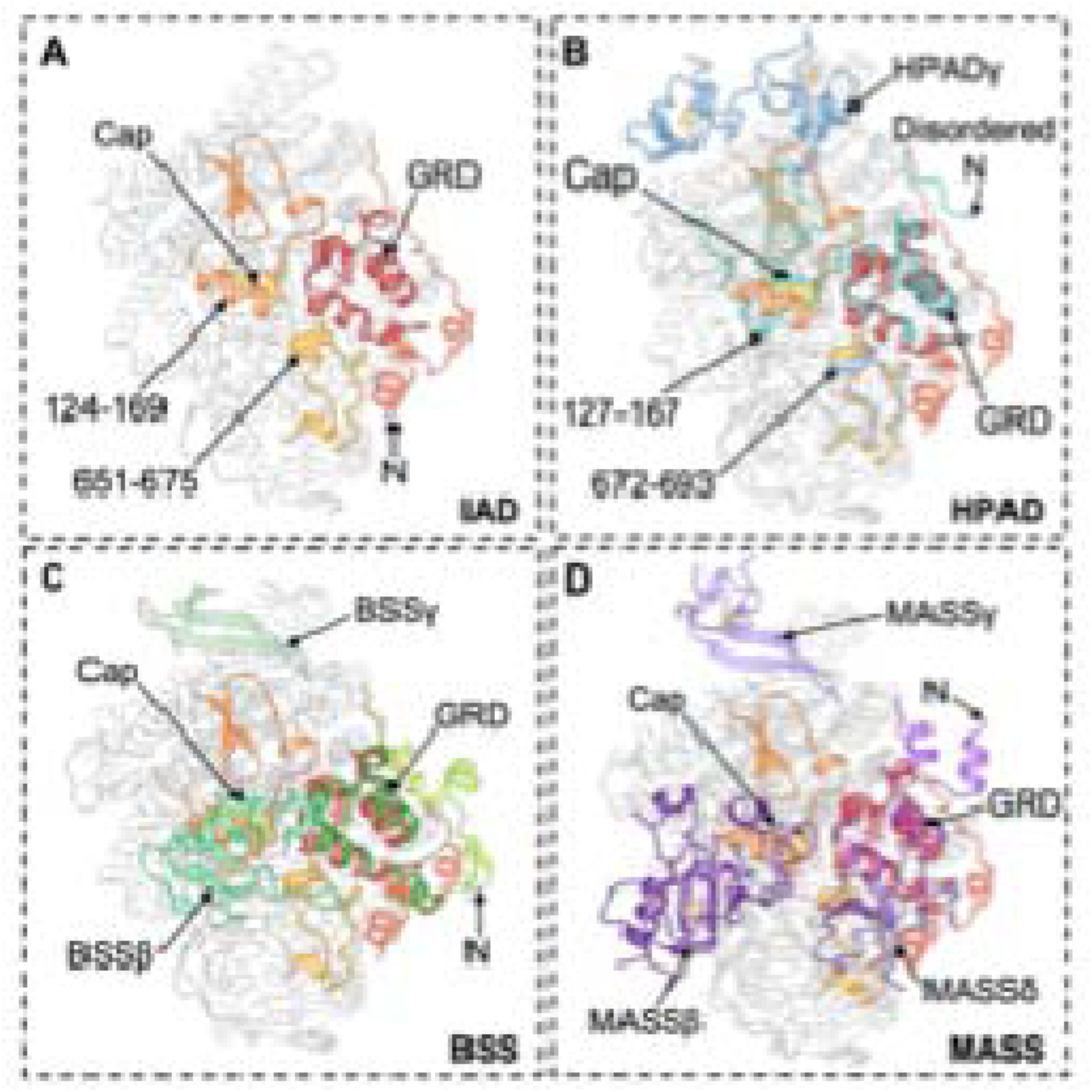
Comparison of *Ou*IAD with HPAD, BSS, and MASS. **A.** Secondary structural features observed surrounding the GRD in IAD. The N-terminus, extension 1 (residues 124-169) and extension 2 (residues 651-675), the cap helix, and the GRD are indicated by arrows. **B.** Overlay of IAD with HPAD shows conservation of the structural features observed in IAD. The N-terminus, the small subunit γ, the cap helix, extension 1 (residues 127-167), extension 2 (residues 672-693), and the GRD are indicated by arrows. **C.** Overlay of IAD with BSS shows overlap of IAD’s extensions with the small subunit β of BSS. The N-terminus, cap helix, subunits β and γ, and the GRD are indicated by arrows. **D.** Overlay of IAD with MASS shows overlap of IAD’s extensions with the small subunits β, γ and δ of MASS. The N-terminus, cap helix, β and δ subunits, and GRD are indicated by arrows.

Interestingly, the locations of IAD’s and HPAD’s extended regions 1 and 2 overlap with the binding sites for BSS’s β subunit and MASS’s β and δ subunits. BSS’s β subunit overlaps with IAD’s extensions 1 and 2 (**Fig 3C**) whereas MASS’s β subunit overlaps with IAD extension 1, and MASS’s δ subunit overlaps with IAD’s extension 2 and N-terminal extension (**Fig 3D**). HPAD, which as mentioned above, shares IAD’s extension 1 and 2, and does not contain equivalent subunits to BSS’s β subunit or MASS’s β or δ subunit. HPAD has one subunit (γ) that occupies a similar location to the γ subunits of BSS and MASS (**Fig 3B-D**). Therefore, it seems that extensions 1 and 2 are mutually exclusive with the presence of a β or δ subunit, and as discussed below, may play similar roles. Notably, the β and δ subunits have been implicated in catalysis/radical installation (20, 24), whereas the γ subunit has been shown to play a role in enzyme solubility (25, 26). In HPAD, the γ subunit covers a hydrophobic patch on the large subunit (**Fig S8A**), and these residues are substituted with primarily hydrophilic residues in IAD (**Fig S8B**).

### Extended regions in IAD and HPAD contact GRD as observed for the small subunits of BSS and MASS

Glycyl radical installation requires that the GRD flip out of the enzyme active site for interaction between the Gly loop and the AE. However, following glycyl radical installation, movement of the GRD is likely restricted to protect the glycyl radical species. Therefore, contacts made to the GRD are of interest. Here, we find that the loop extensions in IAD and HPAD make extensive contacts with their respective GRDs (**Fig 3A-B, Fig S9A-B**). BSSβ, which also makes extensive contacts with BSS’s GRD (**Fig 3C, S9C**), is known to regulate the opening of the GRD for radical installation and subsequent closing of the enzyme for turnover (24). Although the function of MASSβ and δ has not yet been fully characterized, these subunits also make extensive contacts with the MASS GRD (**Fig 3D, S9D**).

Additionally, the N-terminus of IAD wraps around the GRD like a seatbelt, seemingly preventing the GRD from moving outward for radical installation (**Fig 3A, S9A, S10A**). ∼600 of ∼1750 Å^2^ coverage of the IAD protomer by the N-terminus that was mentioned above is to the GRD region. Movement of the N-terminus away from the IAD protein core would expose the GRD and allow it to move (**Fig S10B**). The N-terminus of BSS also makes substantial contacts with the GRD, and the N-terminus of MASS makes a minor contact (**Fig 3C-D, S9C-D**). In contrast, isethionate sulfite lyase (IslA) (27) and hydroxyproline dehydratase (HypD) (28), like most other GREs, have far less extensive contacts between regions of the protein and the GRD, perhaps allowing them to more readily move outward for radical installation (**Fig S9E-F**).

### Loop extensions in IAD and HPAD are located near putative substrate channel as observed for the small subunits of BSS and MASS

In addition to regulating the mobility of the GRDs, the small subunits of BSS and MASS have been proposed to regulate substrate entry/product release (20, 24). GRE active sites are buried, requiring substrate-channels and/or substantial conformational changes for substrate binding and/or product release. A conserved substrate channel for GREs has been identified using published crystal structures and programs such as Caver (**Fig S11A**) (27). In most cases, Caver predicts the channel in the substrate-free state of the enzyme but not when substrate is bound, and the channel is closed. IAD appears to have a similar channel, but in this substrate-bound structure, the channel is closed. IAD’s Trp392, which was predicted by homology modelling to form π-stacking interactions with the substrate I3A indole ring (21), appears instead to be blocking the putative substrate channel in this substrate-bound state (**Fig S11B-C**). In silico removal of the side chain of Trp392 from the model allows Caver 3.0.3 to predict the channel in this substrate-bound state. Experimental substitution of Trp392 with a small amino acid, Ala, has been previously shown to abolish all enzyme activity, whereas substitution with the larger Phe, retained some activity (21). It seems reasonable that another large, hydrophobic residue would still be able to block the substrate channel and allow for turnover, while a smaller residue like Ala would not be sufficient to entirely block the channel and may be more disruptive to catalysis.

In BSS, entrance to the substrate channel is blocked by its β subunit, which, as mentioned above, overlaps with extension 1 of IAD (**Fig 3C, S11E**). A C-terminal helix bundle in BSS was observed to move inward upon binding of fumarate and then moved inward even more upon binding of toluene and the β subunit (**Fig S11D**) (19). Extension 2 of IAD (residues 651-675) extends from a homologous C-terminal helix bundle. MASS β similarly blocks the predicted MASS substrate channel (**Fig S11E**). Additionally, MASS contains a Trp residue (Trp185), positioned similarly to Trp392 in IAD, blocking the substrate channel (**Fig S11E**). These comparisons suggest that GREs may employ small subunits, additional secondary structures, large hydrophobic residues, or a combination of these features to regulate substrate entry or product release from the active site.

### Structure of substrate-bound state of IAD is revealed

Excitingly, all four protomers of the IAD homotetramer have the substrate I3A bound in the active site (**Fig 4A, Fig S12**). Because the enzyme used to prepare the grids did not have the glycyl radical installed, the bound substrate cannot be turned over, allowing for the visualization of a precatalytic state. Overall, the active site of IAD reflects that of HPAD, with a similar substrate binding mode where the carboxylate moiety is oriented toward the catalytic cysteine (Cys500 in *Ou*IAD) (**Fig 4B**). The indole ring of I3A is surrounded by residues Phe401, Phe517, Leu616, Leu730, and Ile732 in a hydrophobic pocket (**Fig 4C**). Of these, Phe401 and Leu616 were identified by homology modelling by the Balskus lab and substitutions of these residues were found to completely abolish enzyme activity (21). Arg226 stacks on top of the indole ring of I3A with an Arg226-NH ^+^ to I3A centroid distance of 3.1 Å, making a putative a cation-π interaction (**Fig 4D**). Arg226 was predicted to make a cation-π interaction, but Trp392 was the amino acid predicted to stack against the indole ring (21). However, as mentioned above, Trp392 does not directly contact I3A and instead appears to gate substrate entry. The only stacking interaction to the indole of substrate is the one made by Arg226, likely explaining why substitution of this residue with Glu, Met, and Lys are catalytically inactive (21).

**Figure 4.**
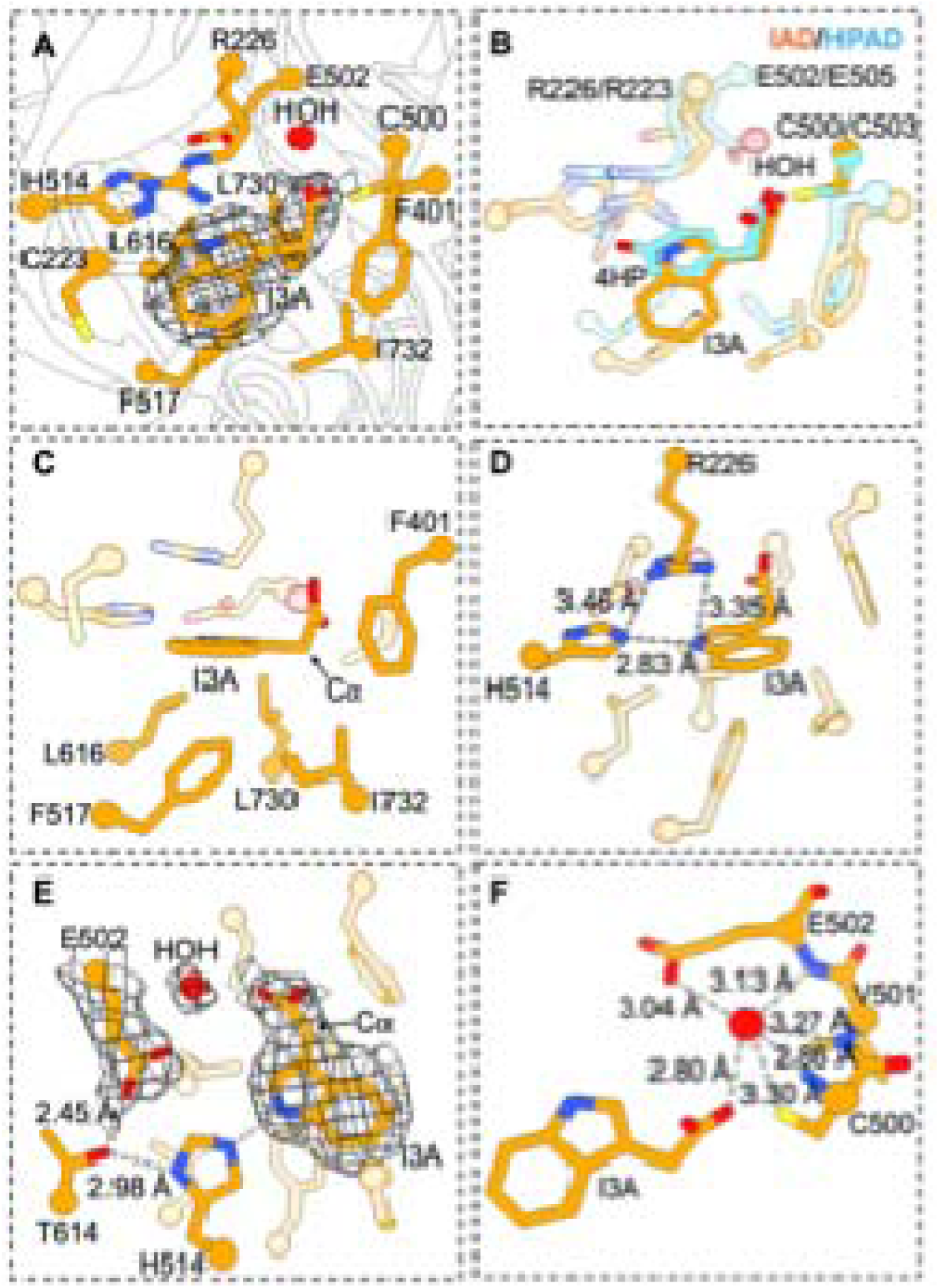
The active site of *Ou*IAD is organized to bind the substrate indole-3-acetate. **A.** Cryo-EM map density for the substrate indole-3-acetate (I3A) in the active site of *Ou*IAD (mesh view, map contoured to r.m.s. level 6.5). **B.** Overlay of IAD and HPAD active sites shows a similar substrate binding mode. **C.** The hydrophobic pocket that surrounds the indole moiety of substrate is composed of F401, F517, L616, L730, and I732. **D.** H514 hydrogen bonds to the indole ring nitrogen of I3A. R226 stacks on the indole ring of substrate in a cation-π interaction and hydrogen bonds to H514. **E.** A water molecule mediates the hydrogen bond network between E502 and I3A. Cryo-EM map density around water molecule, I3A, and E502 contoured to r.m.s. level 6.5 **F.** The water atom observed in the active site is coordinated by I3A, C500, E502, and the backbone nitrogen atoms of the Cys loop. IAD carbons in orange, nitrogens in blue, oxygens in red, sulfur in yellow.

His514 is positioned to hydrogen bond to the indole ring nitrogen of I3A (**Fig 4D-E**). His514 was predicted to be in the active site, but not to be a potential hydrogen bond acceptor of the substrate indole nitrogen nor to be close to Glu502 to potentially affect its pK_a_ (**Fig 4D-E**) (21). Interestingly, substitution of His514 with Ala or Glu has a modest effect on catalysis despite making what appear to be important interactions with substrate (**Fig 4D**). Glu502 comprises a part of a conserved CXE motif where C is the catalytic cysteine. In the structurally characterized GREs that contain this motif, the Glu directly hydrogen bonds to the substrate of the specific GRE and to the backbone nitrogen atoms of the Cys loop **(Fig S13A-D).** We were therefore surprised to see Glu502 of IAD hydrogen bonding to I3A indirectly through a resolved water molecule that also interacts with the backbone nitrogen atoms of the Cys loop (**Fig 4E-F, S12D, S13E**). In other GREs, the Glu carboxylate occupies the position of the water in IAD, interacting with the backbone nitrogen atoms of the Cys loop (**Fig S13F-I**). This water molecule is visualized in the active site of all four protomers of the model and is placed with high condfidence, reporting an average Q-score of 0.909. Fu *et al.* showed that Glu502 is essential for catalysis in IAD and proposed that it might be a crucial proton donor to the ⍰ carbon of substrate concomitant with decarboxylation. However, these structural data show that Glu502 is not positioned well to perform this role (**Fig 4E**). In fact, there is no observable residue or solvent molecule that could easily provide a proton to the ⍰ carbon of I3A (**Fig 4C**). The environment near the ⍰ carbon of I3A is entirely hydrophobic (see below for further discussion).

Overall, the homology modeling was able to predict the identity of many amino acids in the active site. Cys223, which is found at the opposite end of I3A in the active site from the catalytic Cys500 (**Fig 4A**), is an exception. However, the positioning of residues with respect to IAD is different enough from the homology model to warrant a re-investigation of residue roles and of the putative IAD mechanism.

### The active site architecture of IAD is consistent with a Kolbe-type decarboxylation mechanism

Except for HPAD, all GREs are thought to employ the catalytic thiyl radical species to initiate catalysis on a substrate via HAT (12) (**Fig 1A**). Based on both experiment (17) and computation (29), HPAD has been proposed to use the catalytic thiyl radical species to initiate catalysis via electron transfer instead of HAT (**Fig S14A-B**). In this Kolbe-type decarboxylation reaction (**Fig S14A**), electron transfer from the substrate carboxylate to the thiyl radical would generate a substrate acyloxy radical and a cysteine thiolate. This Cys thiolate would be subsequently protonated by an active site residue (**Fig S14A**). In both cases, catalysis would end with HAT from the catalytic cysteine to the product radical species (**Fig S14A-B**).

Although there is no experimental evidence that HPAD’s substrate, 4HP, or IAD’s substrate, I3A, are bound in their deprotonated forms, in the absence of any obvious protein-substrate interaction that would be expected to raise the pK_a_ of substrates’ carboxylates above ∼4, it is generally assumed that both 4HP and I3A are bound as carboxylates and not carboxylic acids. Furthermore, computational studies performed on HPAD support 4HP binding in the active site of the enzyme in the deprotonated, or carboxylate, form rather than as a carboxylic acid (29). If either 4HP or I3A did bind in the carboxylic acid form, hydrogen atom abstraction from the carboxylic acid could initate the reaction. However, such a hydrogen atom abstraction would be expected to be uphill with a bond dissociation energy (BDE) of ∼110 kcal/mol for a carboxylic acid hydrogen versus ∼80-90 kcal/mol for the cysteine thiol.

Both a HAT and Kolbe-type mechanism have been proposed for IAD (**Fig 5A-B**) (21). Each proposal has a role for the thiyl radical at Cys500 and a role as a general acid for Glu502. However, the roles are different and require a different orientation of the amino acid side chain with respect to substrate (**Fig 5A-B**). For a Kolbe-type mechanism to be favored over a HAT mechanism, the catalytic Cys500 would be expected to be positioned very close to the substrate carboxylate. We find the Cys500-Sγ to I3A-carboxylate oxygen distance is very close at 2.44 Å on average in the four chains (**Fig 5C**), closer even than the Cys500-Sγ to I3A-C⍰, which is 4.07 Å on average in the four chains (**Fig 5D**). Notably, both mechanisms involve HAT between a Cys500 thiol and a product radical species, consistent with a ∼4 Å distance (**Fig 5A-B**). The role of Glu502 as a general acid is quite different in the two mechanistic proposals. For the HAT mechanism, Glu502 protonates the substrate C⍰ concomitant with I3A decarboxylation (**Fig 5A**), whereas in the Kolbe mechanistic proposal, Glu502 protonates Cys500 (**Fig 5B**). Here we find that Glu502 is too far (∼6 Å) from the substrate C⍰ to serve as a direct proton donor or an indirect proton donor via the tightly coordinated water molecule, which sits at a distance of ∼5 Å from I3A-Cβ (**Fig 5E**). As mentioned above, there are no other putative general acids near C⍰ of substrate; the environment is completely hydrophobic except for Cys500 (**Fig 4C**). In contrast, Glu502 appears to be perfectly positioned to protonate Cys500 via the water molecule (**Fig 5F**) as would be required for a Kolbe-type mechanism (**Fig 5B**).

**Figure 5.**
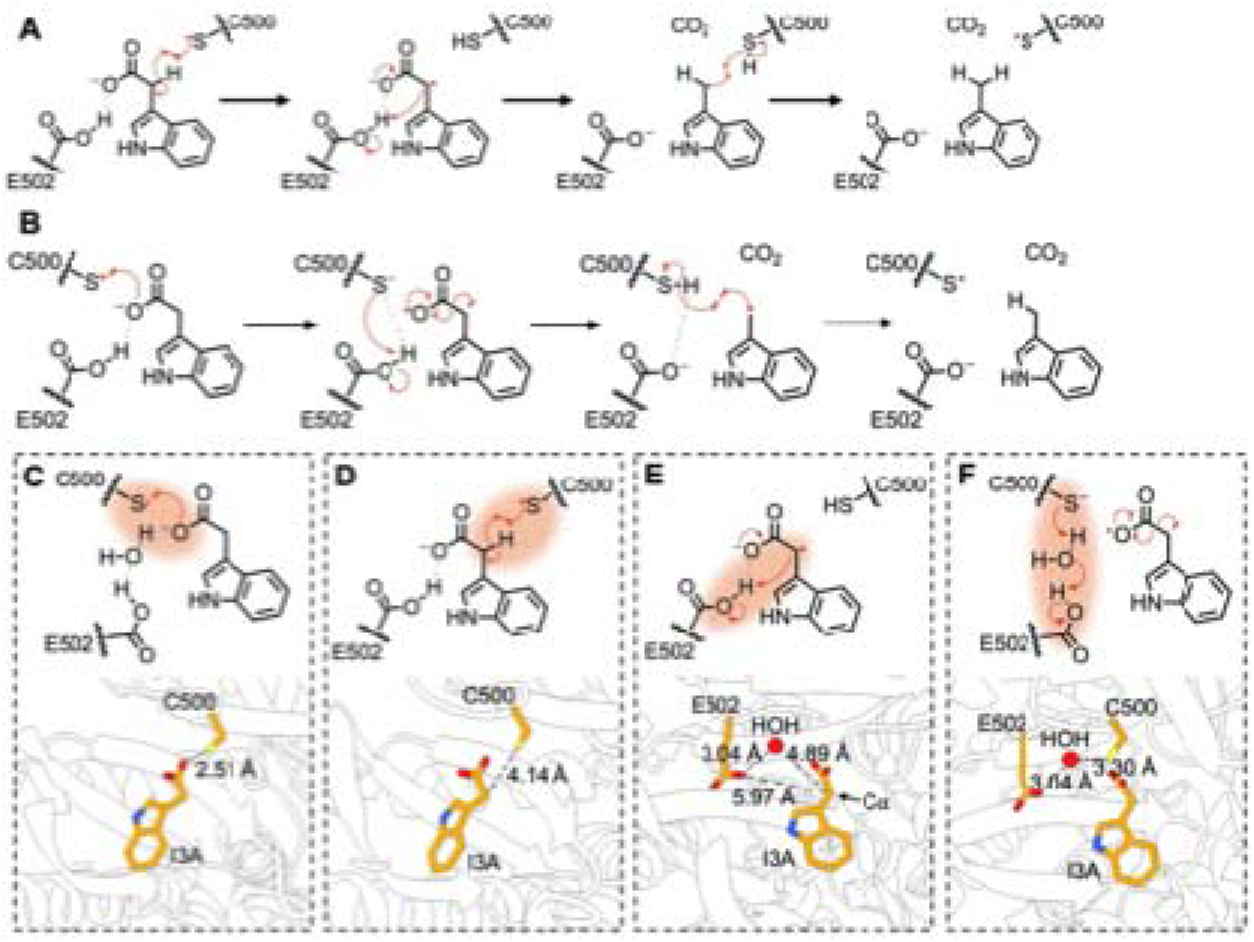
The active site architecture of *Ou*IAD is inconsistent with a hydrogen atom transfer mechanism. **A.** Reaction scheme showing the proposed steps for a hydrogen atom transfer mechanism for IAD. Briefly, the thiyl radical abstracts a hydrogen atom from the ⍰ carbon of I3A. The substrate rapidly decarboxylates and a proton is donated to the ⍰ carbon by Glu502. The product radical can re-abstract the thiol hydrogen atom, reforming the thiyl radical and forming skatole. **B.** Reaction scheme showing the proposed steps for a Kolbe-type decarboxylation for IAD. Briefly, the carboxylate of I3A undergoes one electron oxidation by the thiyl radical. The resulting thiolate is protonated by Glu502 and the substrate radical readily decarboxylates. The product radical re-abstracts a hydrogen atom from Cys500, reforming the thiyl radical and forming skatole. **C.** Step 1 of a Kolbe mechanism requires the thiyl radical to be positioned near the I3A carboxylate. Our structure is consistent with this positioning. **D.** Step 1 of an HAT mechanism requires the thiyl radical to be close to the ⍰ carbon of I3A. Structure shows a 4.14 Å distance. **E.** Step 2 of an HAT mechanism requires a proton donor near the ⍰ carbon of substrate radical. In our structure, C500 and E502 are not positioned well or at appropriate distances to serve these functions. **F.** Step 2 of a Kolbe mechanism requires protonation of the catalytic cysteine thiolate. In our structure, E502 is positioned well to perform this role.

All known structures of GREs to date, including this one, have been captured in the unactivated state (i.e., no glycyl radical has been installed). As such, we cannot rule out the possibility that the active site architecture shifts in some way upon radical installation that would impact the catalytic mechanism. Assuming no significant changes to the active site upon radical installation, the IAD structure is consistent with a Kolbe-type decarboxylation mechanism. Yet, previous deuterium incorporation studies were reported to favor an HAT mechanism (21). Briefly, more than one deuteria were found on the skatole product at the 3’-methyl position, which was easier to explain by a HAT mechanism than a Kolbe-type mechanism (**Fig 5A-B, S15A-B**). However, multiple deuteria can also be explained if the thiyl radical of IAD catalyzes deuterium incorporation on skatole post-turnover. Below, we test this idea by adding skatole to activated IAD in D_2_O buffer.

### The thiyl radical of IAD is capable of 3’-methyl hydrogen exchange on skatole

To test if the thiyl radical of IAD can incorporate deuterium at the 3’-methyl position of skatole (**Fig S16**), we anaerobically prepared the cognate *Ou*IAD activating enzyme and assessed its ability to install glycyl radical on *Ou*IAD. *Ou*IAD-AE purifies with a dark brown color corresponding to 5.39 ± 0.48 Fe per monomer (**Fig S17A-B**). When this IAD-AE is used to activate IAD, we observe the characteristic doublet signal by electron paramagnetic resonance spectroscopy, corresponding to 0.52 radicals per monomer of IAD (**Fig S17C**). Active IAD was capable of catalytic turnover to produce skatole from I3A in H_2_O and D_2_O (**Fig S18A-C**), which was confirmed by Liquid Chromatography coupled with Electrospray Ionization and Quadrupole Time-of-Flight (LC-ESI-QTOF) mass spectrometry. Next, we provided activated IAD with skatole in D_2_O buffer (76.5% D_2_O by volume) and again used LC-ESI-QTOF to look for deuterium incorporation into skatole. Two new peaks appear at m/z= 133.0860 and 134.0920, corresponding to D_1_- and D_2_-skatole (**Fig S18D**). These peaks are absent in control samples where AdoMet or the enzymes are omitted (**Fig S18E-F**), indicating that their production is dependent on activated IAD. Therefore, IAD is indeed capable of exchanging the 3’-methyl hydrogen atoms of its native product, skatole, with solvent. In other words, the presence of multiple deuteria in skatole, as observed previously, do not inform on the enzyme mechanism as exchange can happen post-turnover.

## Discussion

IAD adds to the growing class of GRE decarboxylases that catalyze non-oxidative decarboxylation reactions. The substrates of these GREs are arylacetates; the downstream catabolites of aromatic amino acid fermentation (30). GRE decarboxylases act on these arylacetates to produce the malodorous and uremic, neural, and lung toxins *p*-cresol, toluene, and skatole (31–33). These enzymes are found in the genomes of pathogenic microorganisms, specifically *Clostridial* species such as *C. difficile* and *C. botulinum* (34), and in acetogens. It has been hypothesized that the CO_2_ equivalents produced by decarboxylation could be fed into the Wood-Ljungdahl pathway for the production of acetyl-CoA (35). 3-methylindole, commonly known as skatole, was identified over one hundred years ago as a bacterially derived molecule with a potent and characteristic malodor (1). The deleterious effects of skatole are widespread, ranging from urbanism and odor pollution to human health and food waste. It is consequently desirable to understand the molecular basis of skatole formation, however, until recently our knowledge was restricted by the lack of an identified skatole-producing enzyme. The identification of the GRE IAD in 2018 has thus enabled us to directly probe the structure-based catalytic mechanism by which skatole is produced.

Here we present the first structure of an indoleacetate decarboxylase from the gut and oral microbe *Olsenella uli* in a substrate-bound form and find three features that are unexpected. The first unexpected finding is that IAD is a homotetramer as GREs have long been thought to act as dimers in vivo (36). However, tetrameric assemblies have been seen in the crystal lattices of most GRE structures, and those assemblies match the IAD tetrameric arrangement observed here by cryo-EM. Solution methods, gel filtration and mass photometry, also show IAD tetramer formation, raising the question of whether other GREs can also adopt tetrameric states in solution. Interestingly, the IAD tetrameric interface is comprised of α-helices that directly precede the Cys loop of IAD, perhaps regulating conformational dynamics of the Cys loop. Thus, although the presence of a tetrameric state of IAD was a surprise, structural evidence has hinted at this possibility for some time. Further study will be required to fully understand whether oligomeric state changes in IAD or in other GREs could be a mechanism of activity regulation involving the Cys loop.

The second unexpected finding is just how similar IAD is to the small-subunit containing GREs although it does not have a corresponding small subunit. When studies of HPAD and BSS showed that their common small γ subunit was chiefly responsible for improved solubility, functionally covering a hydrophobic patch on the enzyme (20, 26), the question arose as to whether residue substitutions in the region covered by γ subunit could provide solubility without the need for a small subunit. The structure of IAD provides the answer in the affirmative. Here we find hydrophilic amino acid residues in this region and no requirement of a γ subunit.

In addition to the γ subunit, BSS and MASS both have a β subunit and MASS has an additional δ subunit. The β and δ subunits are located near the GRD and/or the entry point to the substrate channel and are thought to play roles in the regulation of GRD movement for radical installation and/or in substrate channel access (18–20, 24). The rationale behind use of small subunits rather than flexible loops to gate substrate access and/or GRD movement has been enigmatic, and the question arose as to whether some GREs would be found to employ protein extensions rather than standalone subunits. IAD and HPAD structural comparisons show protein extensions in the exact locations of BSS and MASS’s small subunits, providing an affirmative answer to this question. Comparisons of IAD, HPAD, BSS and MASS structures show what appears to be an evolutionary trajectory in terms of small-subunit requirements. MASS represents the GRE that requires the largest number of small subunits known: β, δ, γ. BSS replaces two of MASS’s subunits (β and δ) with one subunit (β), and HPAD replaces BSS’s β subunit with protein extensions but keeps the γ subunit for solubility. Finally, IAD has evolved to be completely independent of small subunits. Both in terms of reaction catalyzed and structure, MASS and BSS are a GRE subclass, and we posit here that both in terms of reaction catalyzed and structure, IAD and HPAD are a GRE subclass.

A primary interest in solving the IAD structure was to evaluate the mechanistic proposals for IAD, which will inform inhibitor design efforts. As mentioned above, GREs typically employ HAT as the initial step in the catalytic cycle. However, the GRE decarboxylase HPAD is proposed to use an alternative mechanism, a Kolbe-type decarboxylation, to catalyze the conversion of 4HP to form *p*-cresol. This proposal was based on the crystal structure of HPAD, solved in 2011 (17), in which the binding mode of the substrate 4HP orients the carboxylate moiety near the catalytic cysteine (Cys503 in HPAD) and is supported by computation (21, 29). Despite catalyzing a very similar decarboxylation reaction, IAD was thought to employ a more traditional HAT mechanism based on both computational analysis and biochemical data (21), and we expected the IAD structure to show differential substrate positioning consistent with a HAT mechanism. However, we found just the opposite; that substrate positioning and the active site of IAD was more consistent with a Kolbe-type mechanism. The agreement between IAD and HPAD in substrate positioning and active site composition was the third surprising result (**Fig S19A-B**).

Over the past three decades, GRE structures have revealed how the canonical 10 stranded β/α-barrel fold allows for customization of active site residues to position a substrate molecule precisely such that the desired hydrogen can be abstracted by the thiyl radical (**Fig S19C-F**). Often substrate-positioning allows for stereospecific C–H bond cleavage (27, 28, 37) and cleavage of a C–H bond that is not the weakest bond in the substrate molecule (28). The active site of IAD stands in contrast to these standard cases, showing that, like HPAD, the carboxylate of I3A is closest to the thiyl radical cysteine and that geometry associated with HAT from the ⍰ carbon is non-ideal (**Fig S19A-B**). Furthermore, the IAD active site shows no catalytic residue capable of protonating the ⍰ carbon concomitant with decarboxylation (**Fig 4C**) as required by a HAT mechanism (**Fig 5A**). Instead, the IAD structure allowed us to envision how IAD could use a Kolbe-type mechanism for I3A decarboxylation (**Fig 6**). The first step, formation of the thiyl radical at Cys500, is followed by a one-electron oxidation of the substrate carboxylate by the thiyl radical. Biologically relevant thiyl radicals have been demonstrated to have reduction potentials ranging from ∼0.9 to 1.33 V (38). Whereas the oxidation potential of aliphatic carboxylates is on the order of ∼1.0 V (39), making it thermodynamically feasible that a thiyl radical can oxidize the substrate carboxylate. Substrate decarboxylation and Cys500 protonation would follow the electron transfer (**Fig 6**). For the former, the acyloxy radical formed by the oxidation would be expected to have a half-life of 10^-10^ to 10^-9^ s (40), and thus should readily decarboxylate forming an alkyl radical. For the latter, a water molecule, which interacts with Glu502, is ideally positioned to protonate Cys500 (**Fig 4F**). A protonated Cys500 can then quench the alkyl radical via HAT (**Fig 6**). With the substrate carboxylate group lost, the ⍰ carbon would be more accessible to Cys500 (**Fig 4F**). In the final steps, the glycyl radical is reformed and product leaves (**Fig 6**).

**Figure 6.**
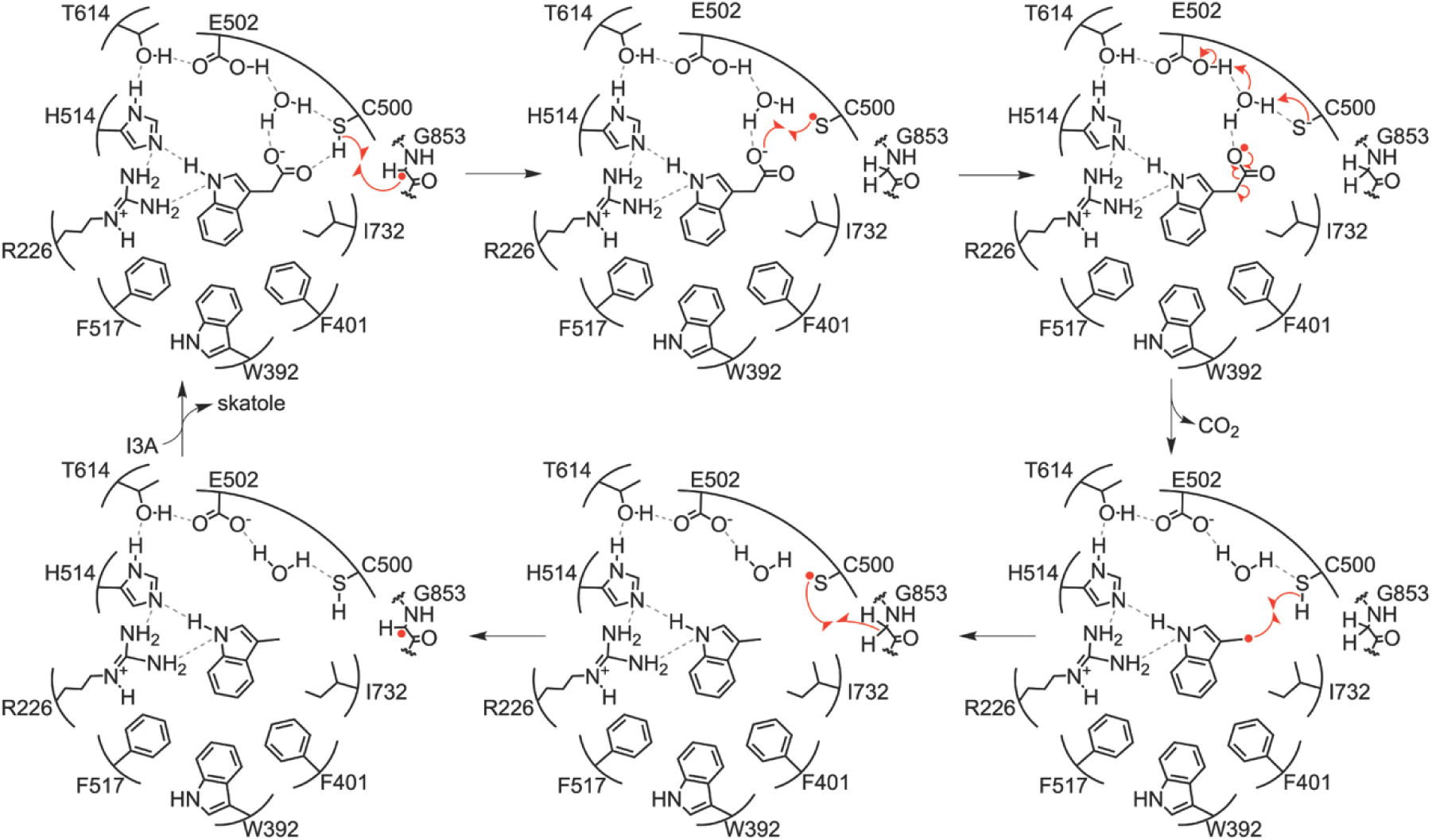
Proposed reaction mechanism for IAD. The glycyl radical at Gly853 abstracts an H atom from the thiol of Cys500, forming a thiyl radical. The thiyl radical then performs a one electron oxidation of the substrate carboxylate, forming an acyloxy radical and a cysteine thiolate at Cys500. The cysteine thiolate is protonated via the water molecule and Glu502 and the acyloxy radical rapidly decomposes to CO_2_ and an alkyl radical. CO_2_ can diffuse out of the active site and the alkyl radical abstracts an H atom from the cysteine thiol at Cys500. Skatole is thereby formed and the radical can return to Gly853 for storage. Skatole then leaves the active site, and the catalytic cycle can begin again.

Prior to this structure determination, experimental data on IAD was either mechanistically agnostic or favored a HAT mechanism (11, 21). These data included: the observation of a minor, but reproducible kinetic isotope effect (KIE); the observation that α,α-methylated-I3A is a partial competitive inhibitor of I3A turnover but is not itself a substrate; and the finding that multiple deuteria can be incorporated at the 3′-methyl carbon of skatole under turnover conditions (21). The KIE data did not strongly favor, nor did it refute, a HAT mechanism. A HAT mechanism in which a C–H bond cleavage is rate limiting would be expected to have a primary KIE in the 3 to 7 range as opposed to a secondary KIE in the 1 to 1.5 range. Measured KIEs for deuterated I3A were in the secondary KIE range ∼1.5 (21), indicating that either C–H bond cleavage does not occur or that the cleavage is not rate limiting. A Kolbe-type mechanism, on the other hand, would be expected to have either no KIE or a secondary KIE, the latter due to the presence of deuterium on the carbon that is also bonded to the carboxylate group that is lost. Therefore, whereas a high KIE would have strongly supported a HAT mechanism, the values obtained were not conclusive.

To examine whether a hydrogen at the I3A ⍰ carbon is necessary for catalysis, Fu *et al.* showed that α,α-methylated-I3A is not a substrate for the enzyme but can partially compete with I3A, suggesting that the lack of turnover is not due to a complete lack of binding (21). Although this result supports a HAT mechanism over a Kolbe-type mechanism, our structure shows hydrophobic residues adjacent to the ⍰ carbon such that methyl substituents would not be easily accommodated (**Fig 4C**). Thus, the lack of turnover of α,α-methylated-I3A can be rationalized by altered binding or active site disruption. Finally, although we find no flaw in the logic that a Kolbe-type mechanism should result in no more than one deuterium in skatole (**Fig S15**), we considered a different possible explanation for the observed multiple deuteria incorporation; that skatole release would be sufficiently slow to allow for exchange of its 3′-methyl protons. Our IAD structure shows that the active site is buried with the putative substrate-channel closed, requiring a protein conformational change for product release. Consistent with a slow product release step, IAD has relatively low turnover numbers: k_cat_ = 2 s^-1^ for *Olsenella scatoligenes* IAD and 11 s^-1^ for *Ou*IAD) (11, 21) compared to HPAD (k_cat_ = 110 s^-1^ for *Clostridioides difficile* HPAD and 135 s^-1^ for *Clostridium scatologenes* HPAD). Our finding in this study that the enzyme can indeed exchange the 3′-methyl hydrogens of skatole with deuteria from D_2_O in an AdoMet- and enzyme-dependent fashion removes this argument in favor of a HAT mechanism. We can now explain the multiple deuteria as being due to a reaction that follows product formation. In addition to explaining the source of multiple deuteria, these data are interesting in their own right. There are multiple examples of GRE reversibility, i.e., GREs acting on their native products. Both BSS and pyruvate-formate lyase (PFL) can catalyze their reverse reactions (41, 42). In fact, the reaction of IAD on skatole is like the forward reaction of BSS, where a proton from the methyl substituent of an aromatic ring system is abstracted by the thiyl radical. In toluene, the bond dissociation energy (BDE) of a methyl C–H bond is approximately 89 kcal/mol whereas in skatole the BDE of a methyl C–H bond is 88 kcal/mol.

Computational studies to probe the basis for the observed deuterium exchange on product as well as the mechanistic proposal put forth in Fig 6 would be useful next steps. As mentioned above a previous computational study carried out after the HPAD structure determination but before the IAD structure determination favored a Kolbe-type mechanism for HPAD but did not support a Kolbe-type mechanism for IAD (21). This study should be repeated using the experimental IAD structure. For HPAD, computational analysis has indicated that deprotonation of the phenol of 4HP by Glu637 would lower the activation energy of the one-electron oxidation (**Fig S14)** such that it becomes more favorable than HAT (21). Additionally, computational analysis suggests 4HP phenolate deprotonation would facilitate the decarboxylation (21), and stabilize the resulting substrate radical (17). In contrast, deprotonation of I3A in IAD was shown to have a minimal effect on carboxylate oxidation and on decarboxylation (21), and although this study was performed on a homology model of IAD, the result is consistent with mutagenesis data showing that the closest residue to the indole N–H, His514, is not essential (21). A recent study on a different family of radical enzymes showed the degree to which an indole moiety can stabilize a radical species via delocalization (43). We are excited to learn what computational analysis can tell us about how the observed IAD active site environment supports skatole formation. We also await further data on phenylacetate decarboxylase (PAD), which lacks the conserved Glu637 observed in HPAD, and can act on both phenylacetate (PA) and 4HP with similar efficiencies (44). PAD is also suggested to use a HAT mechanism (45), more consistent with the majority of GREs, yet in conflict with the mechanism used by HPAD.

Although the work presented here has already expanded our knowledge of the GRE enzyme family considerably, many exciting questions remain. Questions include: what mechanisms are employed by the other members of the GRE decarboxylase subclass and what features may explain any potential difference in mechanism; will other GREs utilize loop extensions as opposed to small-subunits; how many other GREs will be found to employ small-subunits. We hope that future study of GRE decarboxylases will illuminate answers to some of these questions and expand our understanding of the powerful chemistry performed by these enzymes.

## Materials and Methods

Expression and purification of *Ou*IAD and *Ou*IAD-AE are described in the SI appendix, Supplemental Materials and Methods. Briefly, a plasmid containing the gene for *Ou*IAD was expressed in T7 express cells (New England BioLabs). Plasmids containing the genes for *Ou*IAD-AE and sulfur fixation (SUF) were co-expressed in T7 express cells (New England BioLabs). *Ou*IAD and *Ou*IAD-AE contain N-terminal 6xHis tags and were purified aerobically (*Ou*IAD) or anaerobically (*Ou*IAD-AE) via immobilized metal affinity chromatography (IMAC). *Ou*IAD-AE protein was stored anaerobically under liquid nitrogen until use. Iron content in *Ou*IAD-AE was analyzed via Ferene assay (46).

Cryo-EM grids were prepared using the automated SPT Labtech chameleon grid preparation instrument. Quantifoil Active™ grids (300 mesh, hole size 1.2 µm, hole spacing 0.8 µm) were glow discharged at -12 mA for 240 seconds and a final dispense-to-plunge time of 155 ms was used. The cryo-EM data were collected on a Thermo Fisher Titan Krios 300 kV transmission electron microscope equipped with a Gatan G3i K3 camera. The cryo-EM data collected statistics are summarized in **Table S1**. Data processing was performed in RELION 4.0 (47). Reconstruction statistics are summarized in **Table S1**. The final cryo-EM map to 2.45-Å resolution was generated using 330,270 particles. The full workflow, FSC curve, angular distribution plot, and 3DFSC plots are described and shown in the SI Appendix, Supplemental Materials and Methods and **Figs S1-S2**. The local resolution map is shown in **Fig S12**.

An AlphaFold (48) model of *Ou*IAD was docked into the final cryo-EM map using Phenix.dock_in_map (49). A parameter file for I3A (PDB ID: IAC) was generated using eLBOW and bond types, angles, distances, and planes were manually validated. Iterative rounds of model building and refinement of IAD were done using Coot (50) and Phenix.real_space_refine. Phenix.douse was used to place water molecules into the final model. Waters were validated manually and using the check waters function in Coot. Model quality was evaluated using molprobity (51) and EMRinger (52). The refinement and model statistics are summarized in **Table S1**. The final model contains residues 4-878 (of 878) of IAD chains A-D. Figures were created with UCSF ChimeraX (53).

*Ou*IAD activation and turnover assays were performed in a Coy anaerobic chamber under 95% argon, 5% H_2_ atmosphere with <1 ppm O_2_. 50 µM *Ou*IAD, 100 µM *Ou*IAD-AE, 1 mM DTT, 1.5 mM AdoMet, and 100 µM 5-deazariboflavin were illuminated at room temperature using an LED lamp for 2 hr. Turnover assays were performed in buffer containing 76.5% D2O v/v. The mixture was transferred to sample tubes for electron paramagnetic resonance (EPR) spectroscopy, or 2 mM indole-3-acetate or skatole was added and left to react for an additional 2 hr. Reactions were quenched with MeOH and analyzed with LC-MS.

## Data, Materials, and Software Availability

Coordinates and EM data have been deposited in the protein data bank (PDB) (PDB ID: 10AL) (54), the electron microscopy data bank (EMDB) (EMD-75028) (55), and the electron microscopy public image archive (EMPIARs) (EMPIAR-13362) (56). All other data are included in the manuscript and/or the SI.

## Acknowledgments

We thank Dr. Ouliana Pavnova at SPT Labtech for her assistance with cryo-EM grid preparation; Dr. Sarah Sterling and Dr. Jenn Podgorsky at the Cryo-EM facility at MIT.nano for their help with cryo-EM data collection; Dr. Mohan Rajkumar for his help with optimizing the LC/MS method; and Ky Lowenhaupt for her assistance with mass photometry. This work benefited from helpful discussions with Dr. Kelsey R. Miller. This work was supported by NIH Grant 2R35GM126982 (to C.L.D); NIH Grant T32ES007020 (to C.N.I); NSF Graduate Research Fellowship Grant 1122374 (to L.R.F.B); Gilliam Graduate Fellowship (to L.R.F.B); K99/R00 GM145910 (M.C.A.); and the MIT Summer Research Program in Biology. C.L.D. and E.P.B are HHMI Investigators. The Biophysical Instrumentation Facility for the Study of Complex Macromolecular Systems (NSF-0070319) and the Department of Chemistry Instrumentation Facility at MIT are gratefully acknowledged. Specimens were prepared and imaged at the Department of Biology Cryo-EM Facility in MIT.nano. Specimens were screened on a Talos Arctica microscope, which was a gift from the Arnold and Mabel Beckman Foundation. The chameleon was purchased using funds from the HHMI Transformative Technology 2019 Award. Software used in the project was installed and configured by SBGrid.

## Author Contributions

C.N.I., L.R.F.B., M.C.A, and C.L.D. designed research; B.F. and E.P.B. contributed reagents; C.N.I., L.R.F.B., A.P.A., M.C.A, J.C.P., and N.M.G. performed research; C.N.I processed data; C.N.I, L.R.F.B., B.F., E.P.B. and C.L.D. analyzed and interpreted data; C.N.I and C.L.D. wrote the paper. All authors edited and revised the final paper.

## Competing Interest Statement

The authors declare no competing interests.

## Notes

https://www.rcsb.org/structure/10AL

https://www.ebi.ac.uk/emdb/EMD-75028

https://doi.org/10.6019/EMPIAR-13362

### Competing Interest Statement

The authors have declared no competing interest.

### Summary of Updates

Revisions are to correct errors found in proofing.

