## Supporting Information for "The Structural Basis of Malodorant Skatole Formation by the Glycyl Radical Enzyme Indoleacetate Decarboxylase"

**This PDF file includes:**

Supplementary Materials and Methods
Figures S1 to S19
Table S1
SI References

### Supplementary Materials and Methods

#### *Materials*

BL21(DE3) cells (Invitrogen) harboring a pET-28a plasmid with the IAD sequence from *Olsenella uli* strain DSM 7084 (Uniprot: E1QXZ2) or harboring a pET-28a plasmid with the IAD-AE sequence from *Olsenella uli* strain DSM 7084 (Uniprot: E1QXZ4) (1), were kind gifts from the Balskus lab at Harvard University. A plasmid harboring *sufA*BCDSE for iron-sulfur cluster biosynthesis was a kind gift from Tadhg Begley at Texas A&M University. For protein expression and purification, T7 Express cells were purchased from New England Biolabs. Luria broth, 4-(2-hydroxyethyl)-1-piperazineethanesulfonic acid (HEPES, CAS: 7365-45-9), anhydrous imidazole (CAS: 288-32-4), L-cysteine (CAS: 52-90-4), glycerol (CAS:56-81-5), and Fe(II) ammonium sulfate hexahydrate (CAS: 7783-85-9) were purchased from Sigma Aldrich; terrific broth capsules were purchased from RPI; chloramphenicol was purchased from OmniPur (CAS: 56-75-7); benzonase was purchased from EMD Millipore; cOmplete EDTA-free protease inhibitor pellets were purchased from Roche diagnostics; isopropyl β-d-1-thiogalactopyranoside (IPTG, CAS: 367-93-1), lysozyme egg white (CAS: 12650-88-3), kanamycin (CAS: 25389-94-0), were purchased from GoldBio; Talon resin was purchased from Takara Bio; Pierce™ centrifuge columns were purchased from Thermo Scientific; and a Cytiva HiLoad® 16/600 Superdex® 200 pg gel filtration column was used with a Bio-Rad NGC 10 medium-pressure chromatography system. For iron quantitation, Specpure Fe AAS standard solution (CAS: 231-714-2) and sodium meta-bisulfite (CAS: 7681-57-4) were purchased from Alfa Aesar; sodium dodecyl sulfate (CAS: 151-21-3) was purchased from Shelton Scientific; sodium acetate (CAS: 127-09-3) was purchased from Mallinckrodt; L-ascorbic acid (CAS: 134-03-2) and 3-(2-Pyridyl)-5,6-di(2-furyl)-1,2,4-triazine-5′,5″-disulfonic acid (Ferene) disodium salt (CAS: 79551-14-7) were purchased from Fluka Analytical. For cryo-EM grid preparation, indole-3-acetic acid (CAS: 87-51-4) was purchased from Millipore Sigma; dimethyl sulfoxide (CAS: 67-68-5) was purchased from Fisher Scientific and Quantifoil Active grids (Batch:9353) were purchased via the cryo-EM facility at MIT.nano. Glass slides and silicone cassettes for mass photometry were purchased from Refeyn. β-amylase and thyroglobulin standards for mass photometry were purchased from Sigma Aldrich. For in vitro activation and activity assays, *S*-adenosyl-L-methionine (CAS: 86867-01-8) and deuterium oxide (CAS: 7789-20-0) were purchased from Sigma Aldrich; 5'-deazariboflavin (CAS: 19342-73-5) was purchased from SantaCruz Biotechnology; dithiothreitol (CAS: 27565-41-9) was purchased from GoldBio; 3-methylindole (CAS: 83-34-1) was purchased from Thermo Scientific; and sodium bicarbonate (CAS: 144-55-8) was purchased from Mallinckrodt.

#### *Recombinant OuIAD expression*

LB-agar plates containing 50 μg/mL kanamycin were streaked with BL21(DE3) cells containing the pET-28a plasmid encoding the gene for N-terminally 6xHis-tagged *Ou*IAD. A single colony was chosen, grown overnight in LB (Sigma Aldrich) containing 50 μg/mL kanamycin, and 1 mL of the resulting culture was diluted with 1 mL sterile 50% v/v glycerol to create a glycerol stock that was flash frozen in $LN_2$ and stored at -80 °C. A 50 mL LB starter culture with 50 μg/mL kanamycin was inoculated from the glycerol stock and grown for 20 hr at 37 °C with shaking at 220 rpm. LB media (4 L) with 50 μg/mL kanamycin was inoculated with the 50 mL starter culture and then split into 1 L cultures in 2.8 L baffled flasks (ChemGlass) and grown at 37 °C with shaking at 220 rpm until $OD_{600}$=0.6 (approximately 3 hr). Cultures were placed at 4 °C for 30 min without shaking followed by induction of protein expression with sterile-filtered 0.5 mM IPTG and then placed at 17 °C for 20 hr with shaking at 100 rpm. Cells were pelleted by centrifugation (4000 rpm for 15 min), collected in 50 mL falcon tubes, flash frozen in $LN_2$ and stored at -80 °C until lysis.

#### *OuIAD protein purification*

The cell pellet was resuspended in 40 mL lysis buffer (50 mM HEPES pH 7.5, 200 mM NaCl) with the addition of 50 mg lysozyme powder, one EDTA-free protease inhibitor tablet, and 4 μL benzonase. The resuspended cells were lysed with a Branson Digital Sonicator at 65% amplitude for 2 s on and 2 s off for 3 min total and repeated for two cycles. The cell debris was pelleted by centrifugation at 28,000 rcf for 45 min. Supernatant was kept on ice and syringe-filtered with a 0.22

µm filter. A centrifuge column with 2 mL Talon resin (Takara Bio) was equilibrated with 20 column volumes (CV) of lysis buffer and then the filtered cell lysate was applied and allowed to flow through by gravity. The column was washed with 20 CV of wash buffer (50 mM HEPES pH 7.5, 200 mM NaCl, 20 mM imidazole) and protein bound to the column was eluted with 10 CV of elution buffer (50 mM HEPES pH 7.5, 200 mM NaCl, 250 mM imidazole). Eluate was buffer exchanged into IAD SEC buffer (50 mM HEPES pH 7.5, 50 mM NaCl) and concentrated to <2 mL volume using Millipore Amicon centrifugal concentrators (MWCO 50 kDa). A Cytiva HiLoad S200 16/600 gel filtration column was equilibrated overnight using a Bio-Rad NGC FPLC system with 1 CV MilliQ water and 2 CV IAD SEC buffer that were 0.22 µm vacuum-filtered and degassed. Concentrated eluate from the Talon gravity flow column was injected onto the gel filtration column and the column was run at a constant rate of 1 mL/min. Dimer and monomer IAD peaks were separately collected, concentrated with Millipore Amicon centrifugal filters (MWCO 50 kDa), and yield was assessed by absorbance at 280 nm using a NanoDrop™ 2000c spectrophotometer with the following parameters: molecular weight=101.26 kDa and $\varepsilon$/1000=149.66 $M^{-1}$ $cm^{-1}$. The 6xHis tag was not cleaved from the protein for any of the studies shown here. Protein was aliquoted and flash frozen in $LN_2$ for storage at -80 °C.

***Recombinant OuIAD-AE/pSUF expression***

T7 express cells were co-transformed with the pET-28a plasmid encoding the gene for N-terminally 6xHis-tagged OuIAD-AE and the plasmid encoding the genes for sufABCDSE according to the NEB transformation protocol. 200 µL of the transformation was plated on an LB-agar plate containing 50 µg/mL kanamycin and 25 µg/mL chloramphenicol and incubated at 37 °C overnight. A single colony was chosen and grown overnight at 37 °C in LB containing 50 µg/mL kanamycin and 25 µg/mL chloramphenicol. 1 mL of the resulting culture was diluted with 1 mL sterile 50% v/v glycerol to create a glycerol stock that was flash frozen in $LN_2$ and stored at -80 °C. A 50 mL LB starter culture with 50 µg/mL kanamycin and 25 µg/mL chloramphenicol was inoculated from the glycerol stock and grown for 20 hr at 37 °C with shaking at 220 rpm. To TB media (4 L), 50 µg/mL kanamycin, 25 µg/mL chloramphenicol, 47 mg/L L-cysteine, and 150 mg/L Fe(II) ammonium sulfate hexahydrate were added. The TB media was inoculated with the 50 mL starter culture and then split into 1 L cultures in 2.8 L baffled flasks (ChemGlass) and grown at 37 °C with shaking at 220 rpm until $OD_{600}$=1.0 (approximately 4 hr). Cultures were placed at 4 °C for 30 min without shaking followed by induction of protein expression with sterile-filtered 0.5 mM IPTG (GoldBio) and then placed at 20 °C for 20 hr with shaking at 100 rpm. Cells were pelleted by centrifugation (4000 rpm for 15 min), collected in 50 mL falcon tubes, flash frozen in $LN_2$ and stored at -80 °C until lysis.

***OuIAD-AE protein purification***

Protein purification was performed in an Mbraun anaerobic chamber at 4 °C under $N_2$ atmosphere with <1 ppm $O_2$. This protocol was adapted from Vats *et al*. 2025 (2). Lysis buffer, wash buffer, elution buffer, storage buffer, and Talon resin were sparged with argon gas for 30 min and then brought into the anaerobic chamber. The cell pellet was brought into the anaerobic chamber frozen. The cell pellet was resuspended in 40 mL lysis buffer (50 mM HEPES pH 8, 300 mM NaCl, 5% v/v glycerol) with the addition of 50 mg lysozyme powder, one EDTA-free protease inhibitor tablet, and 4 $\mu$L benzonase. The resuspended cells were lysed with a QSonica Q700 Sonicator at 10% amplitude for 2 s on and 2 s off for 6 min total. The cell lysate was transferred to an air-tight 50 mL centrifuge bottle and brought out of the anaerobic chamber. Cell debris was pelleted by centrifugation at 28,000 rcf for 45 min. The centrifuge bottle was then cycled back into the anaerobic chamber. Supernatant was syringe-filtered with a 0.22 µm filter. A centrifuge column with 2 mL Talon resin was equilibrated with 20 CV of lysis buffer and then the filtered cell lysate was applied and allowed to flow through by gravity. The column was washed with 20 CV of wash buffer (50 mM HEPES pH 8, 300 mM NaCl, 5% v/v glycerol, 5 mM imidazole) and protein bound to the column was eluted with 10 CV of elution buffer (50 mM HEPES pH 8, 300 mM NaCl, 5% v/v glycerol, 200 mM imidazole). Eluate was buffer exchanged into IAD-AE storage buffer (50 mM HEPES pH 8, 200 mM NaCl) and concentrated using Millipore Amicon centrifugal concentrators (MWCO 10 kDa). The concentrated protein was a characteristic brown color and a clear protein band near 37 kDa corresponding to IAD-AE was observed by SDS-PAGE (**Fig S17A**). Yield was assessed by absorbance at 280 nm using a NanoDrop™ 2000c spectrophotometer with the following parameters: molecular weight=37.18 kDa and ε/1000=21.89 $M^{-1}$ $cm^{-1}$. The 6xHis tag was not cleaved from the protein for any of the studies shown here. Protein was aliquoted, flash frozen in $LN_2$, and stored in $LN_2$ until use.

***Iron quantitation by ferene assay***

IAD-AE protein concentration was obtained in triplicate using the method described above. An iron quantitation assay was adapted from Abbasi *et al.* 2021 (3). Reagent A was prepared with 1.35 g SDS, 450 µL saturated NaOAc, and 30 mL MilliQ water. Reagent B was prepared with 270 mg ascorbic acid, 9 mg sodium meta-bisulfite, 400 µL saturated NaOAc, and 5.6 mL MilliQ water. Reagent C was prepared with 18 mg ferene disodium salt and 1 mL MilliQ water. 0, 1.66 µM, 3.33 µM, 16.6 µM, 33.3 µM, 50 µM, 66.6 µM, 83.3 µM, and 100 µM Fe standards were prepared in 100 µL volumes using the purchased Fe standard. IAD-AE was diluted to 10 µM in 100 µL volumes in triplicate. Reagent A (100 µL) was added to IAD-AE samples and Fe standards followed by mixing. Reagent B (100 µL) was added to IAD-AE samples and Fe standards followed by mixing. Samples were incubated at 30 °C for 15 min and then reagent C (5 µL) was added to each followed by mixing. Samples were centrifuged at max speed for 5 min, and 250 µL of supernatant was transferred to a Caplugs® Evergreen polystyrene flat bottom 96-well plate. Absorbance at 592 nm was measured using a Molecular Devices SpectraMax® Plus 384 microplate spectrophotometer. A standard curve was constructed and used to assess Fe concentration in the IAD-AE samples (**Fig S17B**).

***Cryo-EM grid preparation***

An unactivated IAD dimer aliquot, i.e., with no glycyl radical installed, was thawed on ice and filtered by centrifugation at 10,000 rcf in a 0.22 µm CoStar™ centrifugal filter (Corning). Concentration was assessed by absorbance at 280 nm, as previously described. Indole-3-acetic acid (I3A) purchased from Millipore Sigma was prepared as a 1 M stock in 100% DMSO and diluted to 100 mM with IAD buffer (50 mM HEPES pH 7.5, 50 mM NaCl) that had been double filtered with a 0.22 µm filter. IAD was diluted to 10 mg/mL (98.76 µM) in the same double filtered IAD buffer and the 100 mM I3A stock was added to the protein solution for a working concentration of 1 mM I3A and 0.1% v/v DMSO immediately prior to cryo-EM grid preparation. A holey carbon-coated copper Quantifoil Active™ cryo-EM grid (300 mesh, hole size 1.2 µm, hole spacing 0.8 µm; batch no:9353) was prepared using the SPT Labtech chameleon automatic cryo-plunging system. The grid was glow discharged at -12 mA for 240 s and two-stripe mode was utilized for grid characterization. A final dispense-to-plunge time of 155 ms was used. The plunge temperature was 24.2 °C; the ethane temperature was -174 °C; shroud humidity was 81%.

***Cryo-EM data collection***

Data collection information is summarized in the data collection table (**Table S1**). Cryo-EM grids were screened, and data was collected at the Cryo-EM facility at MIT.nano. The data used for single-particle analysis was collected on a Thermo Fisher Titan Krios 300 keV transmission electron microscope equipped with a Gatan K3 direct detector, a Gatan BioQuantum image filter, and a Ceta 16M camera. Data collection was automated with EPU software (ThermoFisher Scientific). Collection parameters were as follows: data were collected at 105,000 x magnification (0.822 Å/pixel), 50 frames, 1.04 electrons/$Å^2$/frame dose, and defocus range 0.2 – 2.0 µm. This data set contained 8,927 movies.

***Cryo-EM data processing***

All data processing was performed in RELION 4.0 (4) according to the workflow shown in **Fig S1**. Aberration free image shift (AFIS) groups were created, and the 8,927 movies were patch motion corrected (6 x 6 patches) using a pixel size of 0.822 Å. Contrast transfer function (CTF) estimation was performed using the implementation of CTFFIND-4.1 in the RELION wrapper. Laplacian-of-Gaussian autopicking was done using an 80-160 Å filter on a randomized subset of 100 micrographs. The ~68,000 picked particles were extracted with a box size of 360 and downsampled to a box size of 180 (1.644 Å/pixel). The particles were 2D classified (100 classes) using an EM algorithm with a 160 Å mask and four of the resulting class averages (~52,000 particles) were selected to train a Topaz model for autopicking. The resulting Topaz model was used to pick particles from the entire set of micrographs, and the particles (2.74 million) were extracted with a box size of 360 and downsampled to a box size of 240 (1.233 Å/pixel). Particles were 2D classified in subsets of 500k (200 classes) using a VDAM algorithm and a 200 Å mask. 1.5 million particles were selected and subject to additional 2D classification (100 classes) using an EM algorithm and a 200 Å mask. 1.22 million particles were chosen and combined into a single .star file for initial 3D reconstruction (2 classes) with a 200 Å mask. The overall *ab initio* reconstruction was reported as 6-Å resolution. The better of the two classes (~598k particles) was selected for 3D refinement, resulting in a 3.48-Å resolution map. Subsequent 3D classification resulted in four out of six (~460k particles) classes being retained for 3D refinement which resulted in a map to 3.52-Å resolution. Given the visual symmetry of the map, 3D refinement with C2 symmetry imposed was performed, however, application of symmetry during refinement had no impact on map resolution nor quality. A mask was created using this refined map at a volume threshold of 0.00871 (UCSF ChimeraX version 1.9) (5) with an extension of 6 pixels and softening of 6 pixels. Masked map sharpening in RELION resulted in a map to 3.18-Å resolution. These particles were then re-extracted with a box size of 360 to retain the data collection pixel size (0.822 Å/pixel). Initial 3D reconstruction with this particle stack resulted in a 4.4-Å resolution map. Subsequent 3D refinement produced a map to 3.48-Å resolution. A mask was created using this refined map at a volume threshold of 0.0027 (UCSF ChimeraX version 1.9) with an extension of 6 pixels and softening of 6 pixels. Masked map sharpening in RELION resulted in a map to 3.08-Å resolution. Per-particle CTF refinement was then performed followed by 3D refinement and map sharpening using the previously generated mask, which resulted in a map to 2.64-Å resolution. Particle polishing was performed followed by 3D refinement and map sharpening, which resulted in the final map to 2.45 Å resolution (**Fig S1**). A 3D representation of the angular distribution from the final 3D refinement is shown in (**Fig S2A-B**). RELION reported a range of 2.33-4.36 Å local resolution (**Fig S2A-B**). The Salk Institute for Biological Science's remote 3D FSC server (https://3dfsc.salk.edu/) was used as a secondary assessment of resolution and preferred orientation. 3DFSC reported an average sphericity of 0.939/1 and global model resolution of 2.87-Å resolution (**Fig S2C**).

#### ***Model building and refinement of OuIAD***

Phenix (6) was used for model placement, iterative refinement, and initial water placement. An AlphaFold2 (7) model of *Ou*IAD was used for initial placement into the final EM map using phenix.dock_in_map. The model placement was manually validated, followed by refinement using phenix.real_space_refine. Briefly, simulated annealing and morphing (once each) were followed by 5 macrocyles of rigid body refinement, model regularization, energy minimization, and refining atomic displacement factors. Model building was done in Coot (version 0.8.9.2 EL) (8). A parameter file for I3A (PDB ID: IAC) was generated using eLBOW and bond types, angles, distances, and planes were manually validated in the text file. I3A was placed in all four protomers of the model followed by refinement. Iterative rounds of refinement in Phenix and model building in Coot were performed until model statistics were satisfactory. All four chains (A-D) of the model contain residues 4-878 (of 878). We were not able to visualize the 6x-His tag that was not removed from the protein prior to structure determination. To our knowledge, there is no evidence that 6x-His tags interfere with the structure or activity of GREs. Four substrate molecules (out of four possible) are present in the active sites of chains A-D. Phenix.douse was used to place water molecules into the finalized model of *Ou*IAD with I3A bound in all four protomer active sites. The check waters function in Coot was used to remove waters that: 1) did not have a corresponding map threshold of 1.5 rmsd or higher; 2) had a contact closer than 2.3 Å distance; or 3) had no contacts within 3.5 Å distance. Water molecules were then manually validated. Those that did not have clear, generous spherical density at 1.5 rmsd, were not hydrogen bonded to the protein or other water molecules, or caused clashes with the protein model were removed. The total number of water molecules is 686. Q-scoring was performed in UCSF ChimeraX (version 1.9). The overall average model Q-score is 0.81; the average Q-score for water molecules is 0.894; the average Q score for the water molecule identified in the active site is 0.909; the average Q-score for IAC is 0.849; the average Q-score for protein is 0.814. Model visualization was also performed in UCSF ChimeraX. The CAVER 3.0.3 (9) plug-in in PyMOL (version 2.4.2) was used to visualize the predicted substrate channel in IAD with a probe radius of 0.9 Å.

***Mass Photometry and analytical SEC***

Mass photometry measurements were collected with a Refeyn Two$^{MP}$ mass photometer. A mixture of 0.5 µM β-amylase and 0.15 µM thyroglobulin diluted in *Ou*IAD buffer (50 mM HEPES pH 7.5, 50 mM NaCl) was used as a mass standard. This mixture was diluted 1:9 in-drop on the mass photometer and the resulting mass peaks were used to create a 4-point calibration curve. Appreciable peaks for β-amylase monomer (56 kDa), dimer (112 kDa), and tetramer (224 kDa), and thyroglobulin dimer (670 kDa) were present. The resulting calibration curve had an $R^2$ value of 1.00 and a maximum mass error of 4.7%. *Ou*IAD was thawed from storage at -80 °C and 0.22 µm filtered with a CoStar™ centrifugal filter (Corning). Concentration was assessed by absorbance at 280 nm using a NanoDrop 2000c spectrophotometer (Thermo Scientific) with the following parameters: molecular weight=101.26 kDa and ε/1000=149.66 $M^{-1}$ $cm^{-1}$. *Ou*IAD was prepared at concentrations of 250, 500, 1000, and 2000 nM in *Ou*IAD buffer with a 1:1 molar ratio of I3A. I3A was prepared as a 1 M stock in 100% DMSO, diluted to 100 mM in *Ou*IAD buffer (10% v/v DMSO) and then subsequently diluted into the final protein solutions. Solutions were diluted 1:9 in-drop on the mass photometer such that the *Ou*IAD concentrations for data collection were 25, 50, 100, and 200 nM. Data were collected in duplicate. All movies were collected for 60s. Two$^{MP}$ Discover software was used to analyze data and make figures.

To assess the oligomeric state profile of IAD, a Cytiva Superdex® 200 Increase 10/300 GL gel filtration column was used with a Bio-Rad NGC 10 medium-pressure chromatography system. Following column equilibration with 1 column volume (CV) of MilliQ water followed by 2 CV IAD SEC buffer (50 mM HEPES pH 7.5, 50 mM NaCl), IAD (250 µL, ~1 mg/mL concentration) was applied to the column and run at a flow rate of 0.25 mL/min. Progress of the run was monitored by absorbance at 280 nm.

***OuIAD activation assay***

IAD activation assays were performed in a Coy anaerobic chamber under 95% argon, 5% $H_2$ atmosphere with <1 ppm $O_2$. 5'-deazariboflavin, a photoinducible reduction agent, was used to reduce the active site cluster of IAD-AE to initiate glycyl radical installation. IAD activation buffer (50 mM HEPES pH 8, 200 mM NaCl) was sparged with argon gas for 30 min and cycled into the anaerobic chamber. AdoMet purchased from Sigma Aldrich was prepared as a 50 mM stock in activation buffer. DTT was prepared as a 100 mM stock in activation buffer. 5'-deazariboflavin was prepared as a 2.5 mM stock in DMSO. IAD, IAD-AE, AdoMet, DTT, and 5'-deazariboflavin were cycled into the anaerobic chamber frozen on cold beads. 50 µM IAD, 100 µM IAD-AE, 1 mM DTT, 1.5 mM AdoMet, and 100 µM 5'-deazariboflavin were illuminated at room temperature using an LED lamp for 2 hrs. The activation mixture was then transferred to EPR tubes and frozen in $LN_2$ inside the anaerobic chamber. $O_2$ levels were at no point elevated above 3 ppm. Radical installation was then assessed by spin quantitation using electron paramagnetic resonance spectroscopy.

***Electron paramagnetic resonance spectroscopy of OuIAD***

EPR spectra were collected using a Bruker EMX-Plus spectrometer equipped with a ER4119HS resonator. Cryogenic measurements were conducted using a Bruker/ColdEdge 4K waveguide cryogen-free cryostat. Sample temperature was regulated with an Oxford Instruments MercuryiTC temperature controller. Signals were compared and quantified via double integration relative to a 1 mM Cu(II)EDTA spin standard, which was prepared from a copper atomic absorption standard solution purchased from Sigma-Aldrich. The following general instrument parameters were used unless stated otherwise: 9.36 GHz microwave frequency, 28 ms conversion time, 100 kHz modulation frequency, and 3 G modulation amplitude. Glycyl radical spectra were acquired at a center field of 3350 G, sweep width of 200 G, and a 30 s sweep time, wherein the reported spectra are the result of a 10-scan signal average. Samples were examined at specific temperatures and applied microwave powers to satisfy non-saturating resonance conditions. EPR signal quantifications were accomplished using the software SpinCount (10) developed by Prof. Michael Hendrich at Carnegie Mellon University.

***OuIAD indole-3-acetate and skatole activity assay in $D_2O$***

IAD activation was performed as described above. 1 M HEPES pH 8, 4 M NaCl, $D_2O$, EtOH, and DMSO were sparged with argon gas for 30 min and then cycled into the anaerobic chamber.

Activation buffer (50 mM HEPES pH 8, 200 mM NaCl) was prepared as a 90% v/v $D_2O$ solution by volume using 1 M HEPES pH 8 and 4 M NaCl stocks in $H_2O$. AdoMet was prepared as a 50 mM stock in $D_2O$ activation buffer. DTT was prepared as a 100 mM stock in $D_2O$ activation buffer. 5'-deazariboflavin was prepared as a 2.5 mM stock in DMSO. I3A was prepared as a 100 mM stock in DMSO. 3-methylindole (skatole) purchased from Thermo Scientific was prepared as a 100 mM stock in EtOH. IAD, IAD-AE, 5'-deazariboflavin were cycled into the anaerobic chamber frozen on cold beads. DTT, AdoMet, I3A, and 3MI were cycled in as solids, and solutions were prepared in the anaerobic chamber. After activation, 2 mM I3A or 2 mM 3MI was added and left to react for 2 hrs. Final reactions were 76.5% v/v $D_2O$. Reactions were quenched with two volumes of MeOH. Protein was removed by filtration using 0.22 µm CoStar centrifugal filters. Solutions were transferred to 150 µL high-recovery glass inserts in 2 mL vials (Agilent) for LC-MS analysis. For the no AdoMet control, AdoMet was omitted from the reaction. For the no enzyme control, IAD and IAD-AE were omitted from the reaction.

***Liquid chromatography/mass spectrometry analysis of IAD activity***

Native IAD turnover of indoleacetate and IAD activity on skatole was assessed using Q-TOF LC/MS (Agilent 6545 mass spectrometer coupled to an Agilent Infinity 1260 liquid chromatography system). An Agilent Poroshell 120 EC-C18 (2.7 µm, 2.1 x 50 mm) reverse phase column was used. Solvent A was $H_2O$ with 0.1% v/v formic acid and solvent B was acetonitrile with 0.1% v/v formic acid. The column was maintained at 25 °C. The flow rate was 0.3 mL/min. The LC method was as follows: 0-1 min, 10% B; 1-4 min, gradient from 10 to 100% B; 4-7 min, 100% B; 7-8 min, gradient from 100 to 10% B. The retention time of 3-methylindole was 4.434 min, the retention time of indole-3-acetate was 3.172 min. The LC method described was used with the MS in positive ion mode. A dual AJS ESI source was used with the following parameters: nozzle voltage 1 kV; capillary voltage 3.5 kV; gas temp 300 °C; drying gas 8 L/min; nebulizer 30 psi; sheath gas temp 350 °C; and sheath gas flow 8 L/min. MassHunter software was used for all data processing and visualization.

### Figures

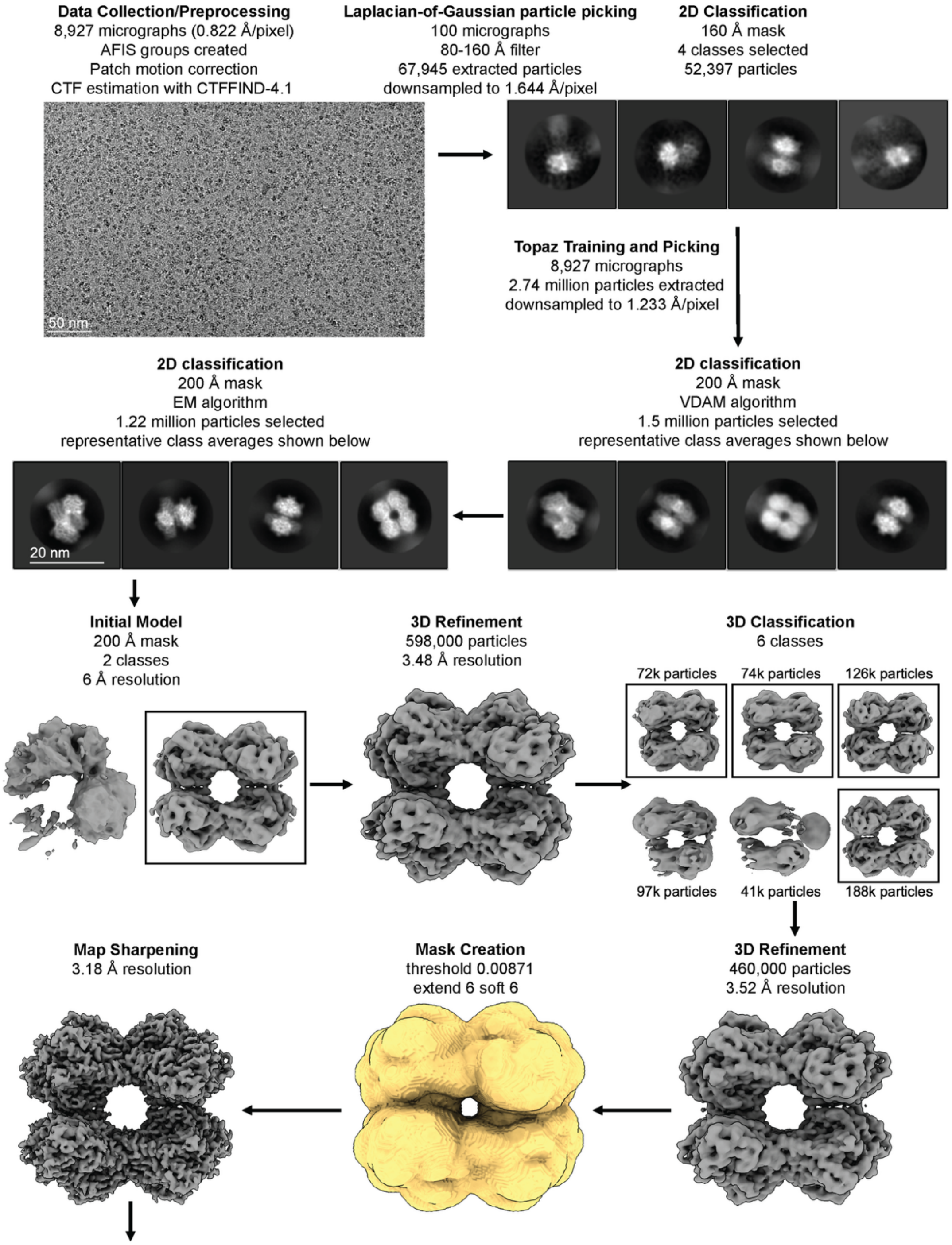
Data Collection/Preprocessing
8,927 micrographs (0.822 Å/pixel)
AFIS groups created
Patch motion correction
CTF estimation with CTFFIND-4.1
50 nm
Laplacian-of-Gaussian particle picking
100 micrographs
80-160 Å filter
67,945 extracted particles
downsampled to 1.644 Å/pixel
2D Classification
160 Å mask
4 classes selected
52,397 particles
Topaz Training and Picking
8,927 micrographs
2.74 million particles extracted
downsampled to 1.233 Å/pixel
2D classification
200 Å mask
EM algorithm
1.22 million particles selected
representative class averages shown below
2D classification
200 Å mask
VDAM algorithm
1.5 million particles selected
representative class averages shown below
20 nm
Initial Model
200 Å mask
2 classes
6 Å resolution
3D Refinement
598,000 particles
3.48 Å resolution
3D Classification
6 classes
72k particles
74k particles
126k particles
97k particles
41k particles
188k particles
Map Sharpening
3.18 Å resolution
Mask Creation
threshold 0.00871
extend 6 soft 6
3D Refinement
460,000 particles
3.52 Å resolution

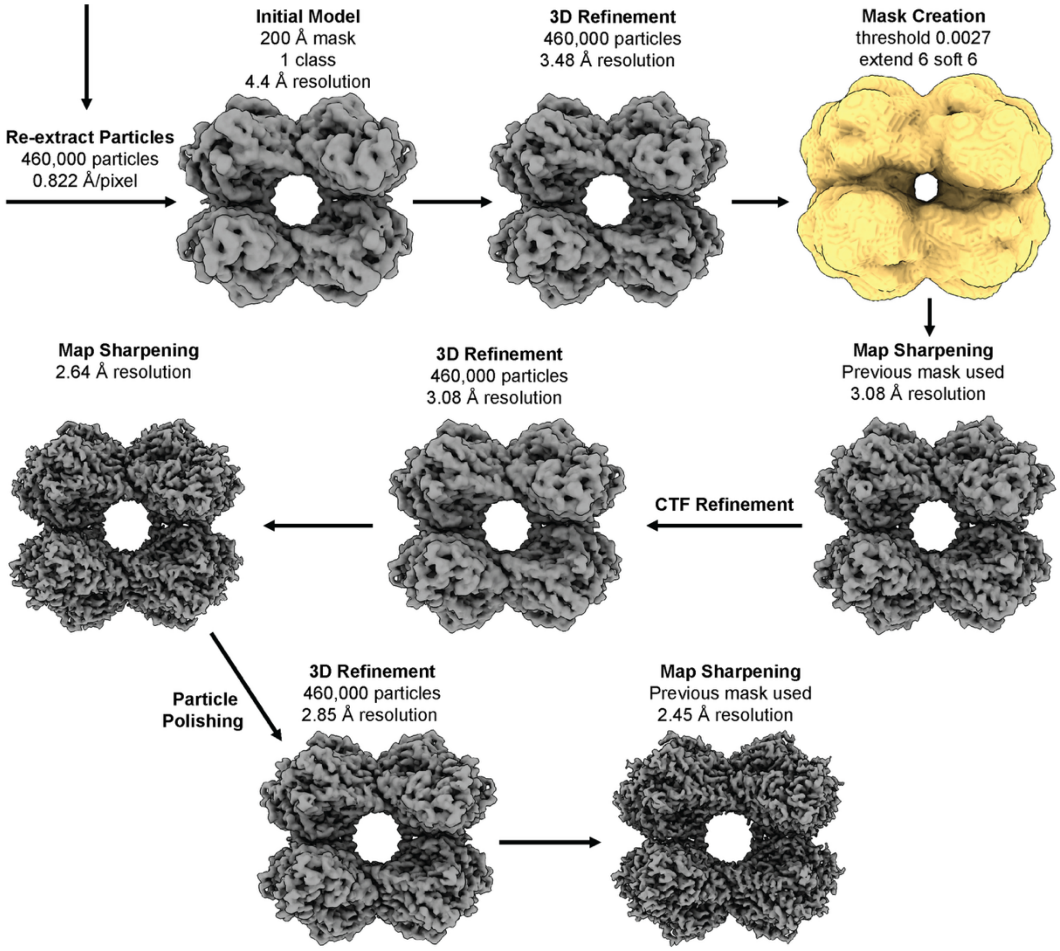


**Figure S1. Cryo-EM data processing pipeline for *Ou*IAD in RELION 4.0.** Scheme describing the processing steps employed to arrive at the final 2.45-Å resolution IAD cryo-EM map. All maps are contoured to 4.5 rmsd.

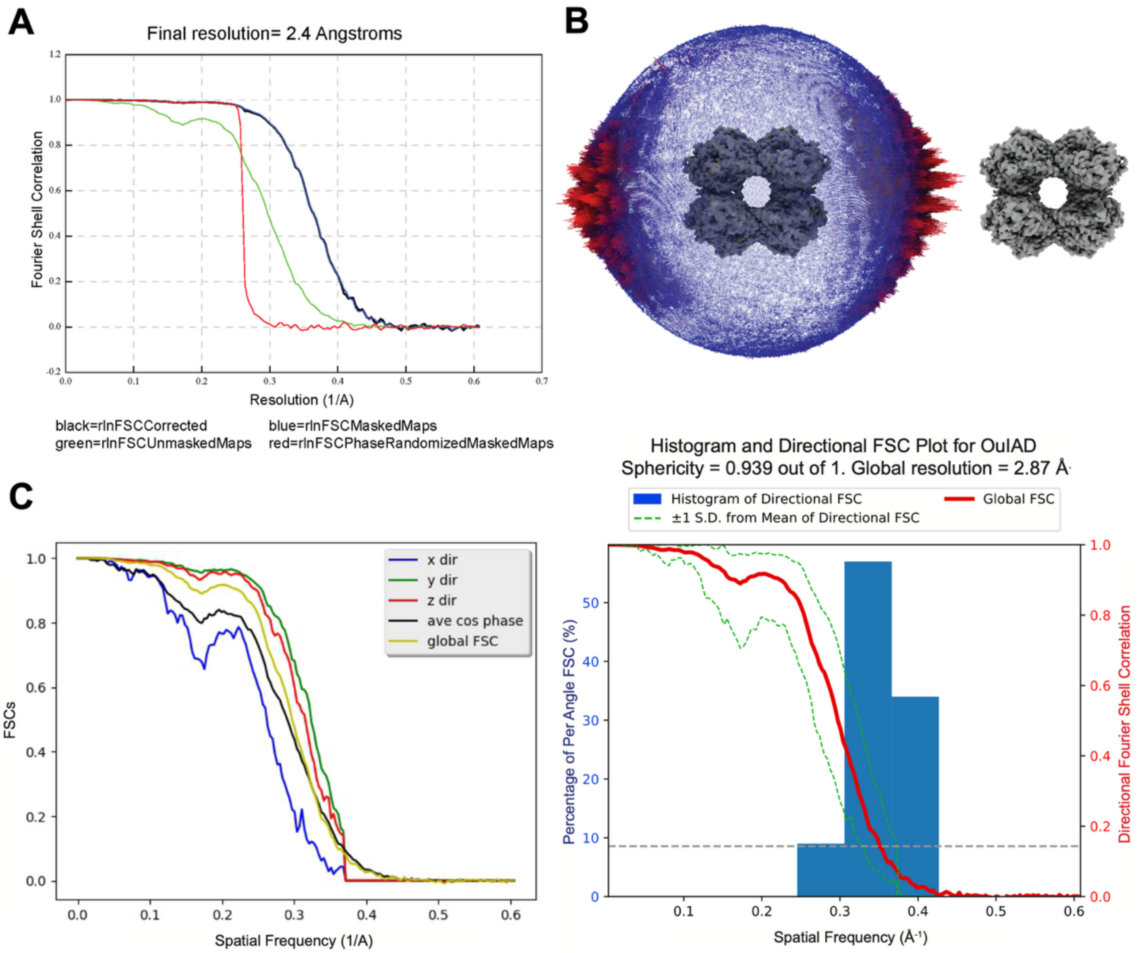


**Figure S2. Fourier Shell Correlation (FSC) curves and angular distribution of the final *Ou*IAD map. A.** FSC curve and final resolution from RELION-4.0. **B.** Angular distribution of final IAD map. **C**. 3D FSC calculation using the Salk Institute for Biological Sciences remote 3D FSC server.

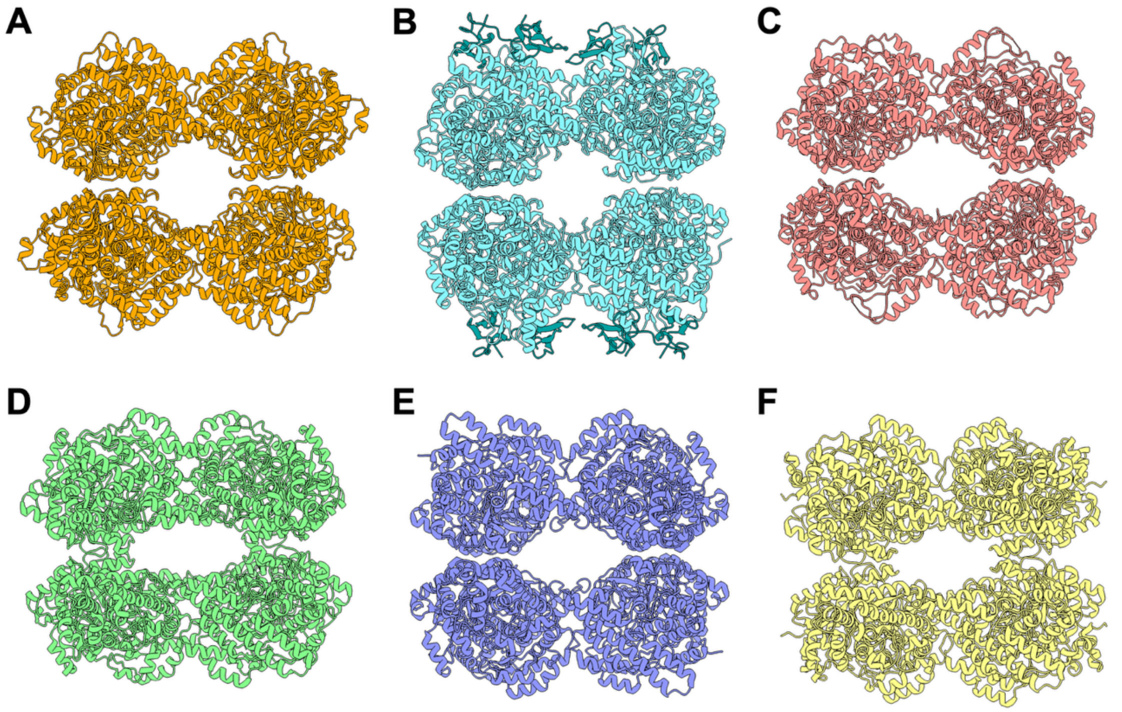


**Figure S3. A GRE tetramer has been observed in the crystal lattice of GREs that have been solved by X-ray crystallography. A.** Indoleacetate decarboxylase (IAD) *this work.* **B.** 4-hydroxyphenylacetate decarboxylase (HPAD) PDB ID: 2YAJ. **C.** Arylacetate decarboxylase (AAD) PDB ID: 7E7L. **D.** Choline trimethylamine lyase (CutC) PDB ID: 5FAU. **E.** Hydroxyproline dehydratase (HypD) PDB ID: 6VXE. **F.** Isethionate sulfite-lyase (IslA) PDB ID: 7KQ3.

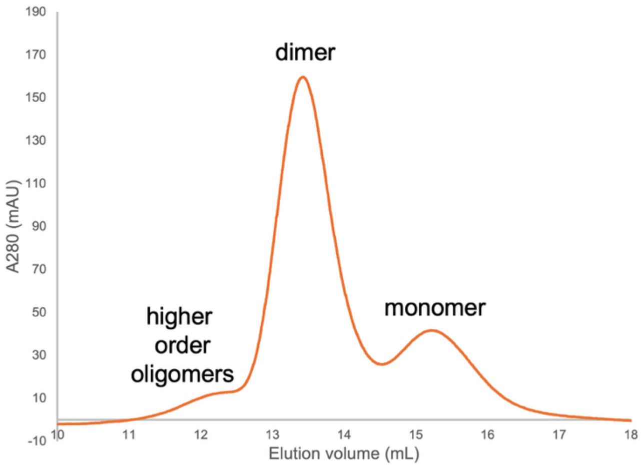


**Figure S4. *Ou*IAD forms higher order oligomers that are observable by analytical SEC.** Size exclusion chromatography (250 µL injection of IAD at 1 mg/mL) shows IAD primarily as a dimer and also as a monomer and higher order oligomer.

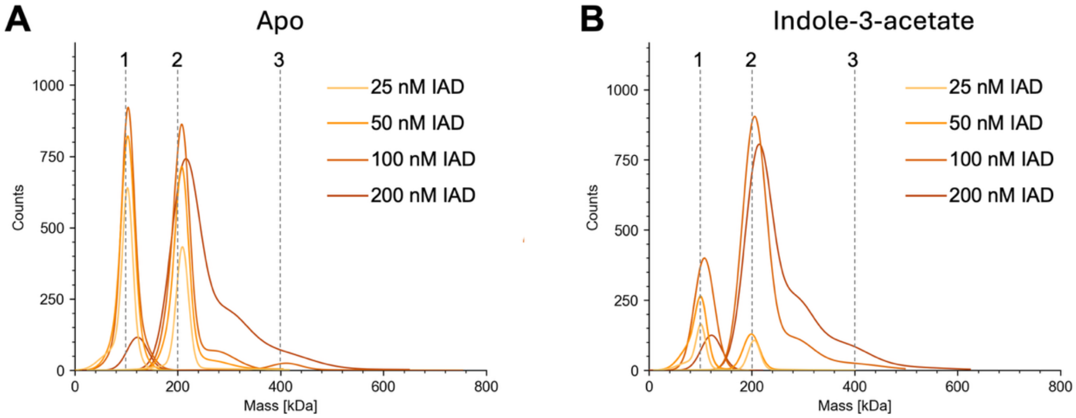


**Figure S5. The *Ou*IAD tetramer can be observed in solution by mass photometry. A.** Increasing protein concentration decreases the population of monomer peak at 100 kDa and leads to the appearance of a tetramer peak at 400 kDa. A dimeric species at 200 kDa is present at all concentration. **B.** Same as in A with the addition of non-saturating amounts of the substrate I3A (1 molar equivalent of I3A in each sample). Peak 1 corresponds to monomer; peak 2 to dimer; and peak 3 to tetramer.

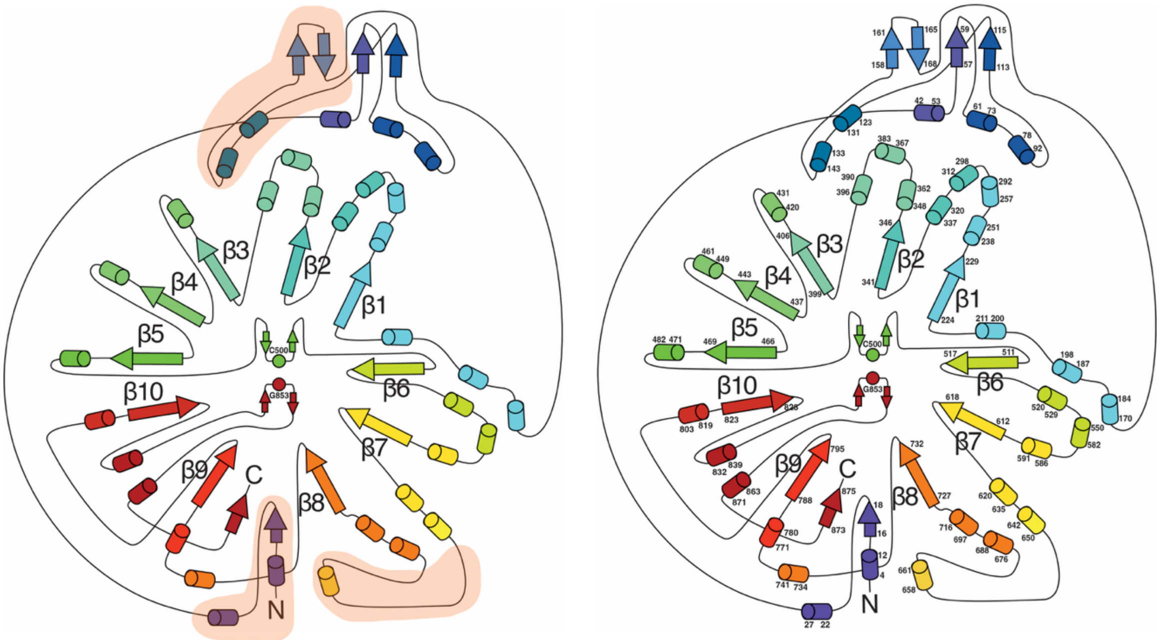


**Figure S6. Topology diagram of *Ou*IAD. Left:** Topology diagram of *Ou*IAD showing the 10-stranded β/α barrel with the N-terminus extension, loop extension 1 and loop extension 2 indicated by orange highlighting. β-strands are numbered in order from the N- to C-terminus. Cys500 of the Cys loop and Gly853 of the Gly loop are labeled. **Right:** Same topology diagram, but with additional residue numbers shown. The cap helix that covers the active site is found between β2 and β3 and is comprised of residues 390-396. The GRD is at the C-terminus of the barrel and is comprised of residues ~832-875.

**A**

```
AAD       SNMLDYKKYEG-QTRGELIWSRLNPLRHT----------------------------AKM  31
PAD       ------MSTQVSQHAPKAPEQMPRKIKLNFD------PNGKMSDRFKKEKEKLFAAPARL  48
IAD       --MEESLVLEMLQTGKTATWPAKNQLQESGEVVDKEVKGTPSTDRTRRMKERFMNAKCKM  58
HPAD      MNVKETKLEDVLKSRGI-------DMKDAYNISEADIP--EAKESTQKLMDIYYTLKVTA  51
                   :  :            ::
```

**B**

```
AAD       DLAASWV----------EDYIDKYGW----------KGSGTFYEFT------------GP  119
PAD       WRDSYWI-------EDDKEFALQRGVHK-----------------------------TTE  132
IAD       QFATDFMISFLTQAEDQKEEA--------KLYSVEGKDEAHTVEEEGWT-KVGQLFSIRE  169
HPAD      WVCASF---FNAQAEALMNEVDAPAENEADSVSVVGAGGGNVTESYGNVISIAKKFGMRK  168
             : :            :
```

**C**

```
AAD       NLYVLKNQIFVDNPICSKETLLDAIHKNW-----------------------EGYEDLQA  614
PAD       CLAAMKKLIFDDKSV-SMEKMVAAITANF-----------------------VGYEDVQN  644
IAD       GLAAIKKLVFEDHKY-TWDEFKEAMDNNFGFVLGAEKGNFSMLNQEIDPEKHMKYAHIHR  684
HPAD      SLSSIKKNVFDDSKF-TIEEMTDAMLNNFGFKTAYETEVFSPDFR-ESTDKSTKYEKIFA  704
           *  :*: :* *    : : :  *:  *:                         * .:
```

**Figure S7. Multiple sequence alignment of known GRE decarboxylases. A.** Alignment of the N-termini of the GRE decarboxylases shows that HPAD and PAD have extended N-termini, similar to that observed in IAD, while AAD has a slightly shorter N-terminus. **B.** Alignment of IAD's extension 1 (residues 124-169) shows that while HPAD has a similar extension, AAD and PAD have only modest extensions in this region. **C.** Alignment of IAD's extension 2 (residues 650-675) shows that HPAD has a similar extension, while AAD and PAD do not. Clustal2 coloring is used.

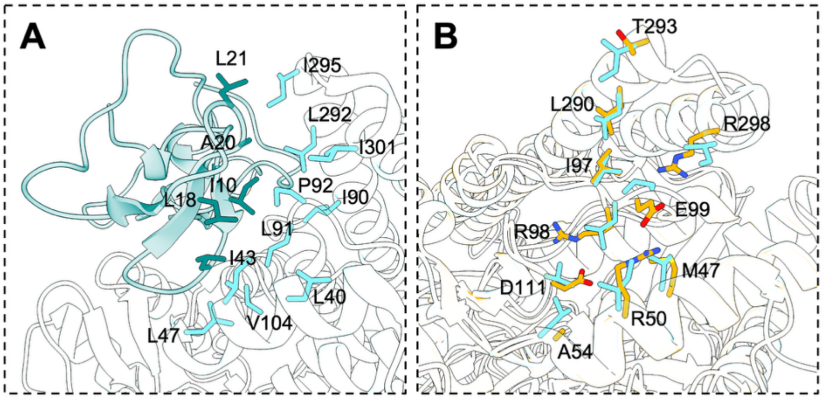


**Figure S8. A hydrophobic patch covered by HPADγ is substituted with hydrophilic residues in IAD. A.** The hydrophobic patch at the interface of HPADα and HPADγ. **B.** In IAD, the residues that comprise the hydrophobic patch on HPADα are substituted with more hydrophilic residues. Only IAD numbering is shown.

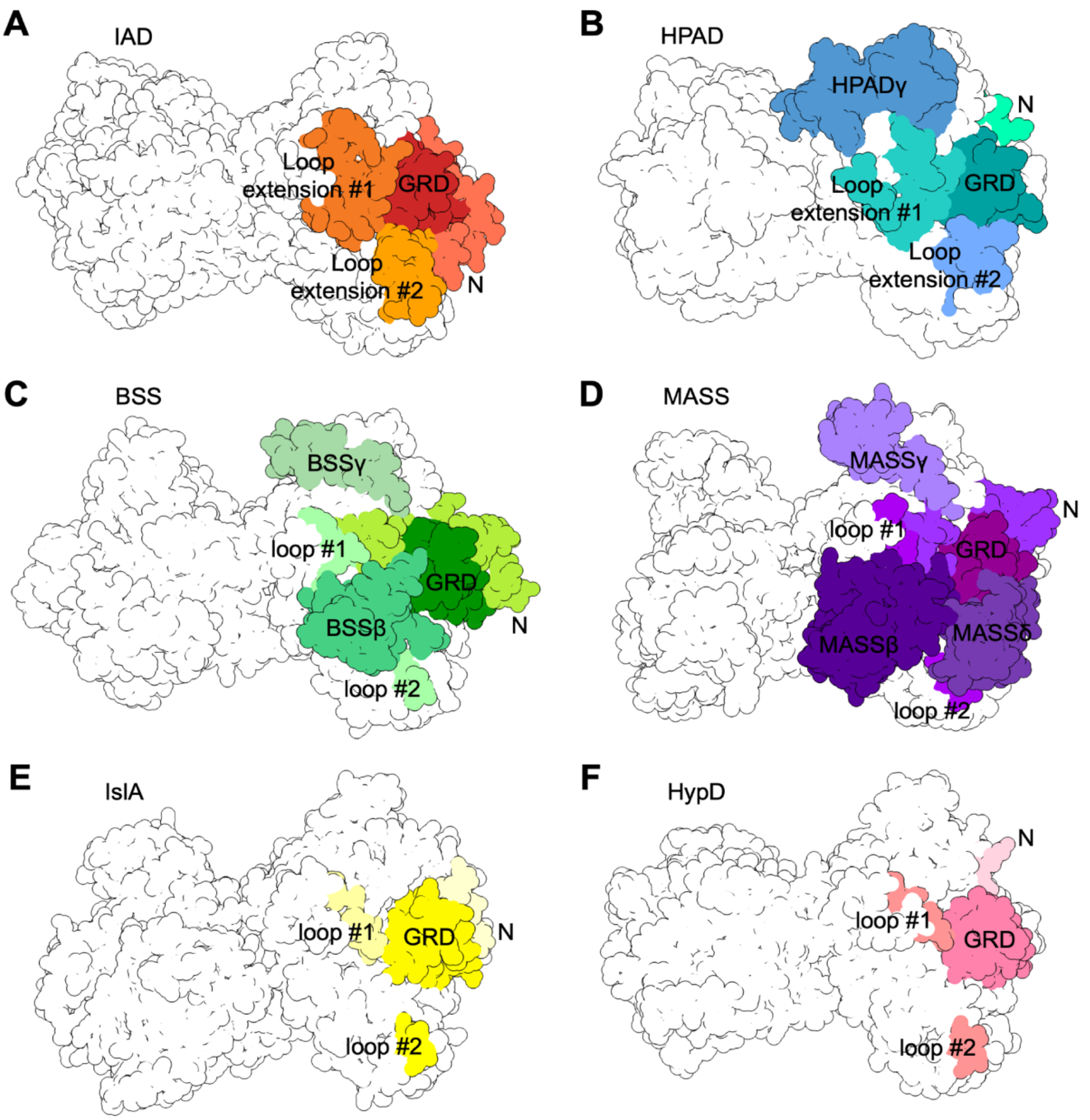


**Figure S9. The glycyl radical domain in GRE decarboxylases and XSSs are more protected. Atom representation of: A.** IAD with GRD, N-terminus, and extensions #1 and #2 colored. **B.** HPAD with GRD, N-terminus, extensions #1 and #2, and γ colored. **C.** BSS with GRD, N-terminus, β, γ, loop #1 and loop #2 colored. **D.** MASS with GRD, N-terminus, β, γ, δ, loop #1 and loop #2 colored. **E.** IslA with GRD, N-terminus, loop #1 and loop #2 colored. **F.** HypD with GRD, N-terminus, loop #1 and loop #2 colored. Loops #1 and #2 in BSS, MASS, IslA, and HypD are the site of the loop extensions observed in IAD and HPAD. N-termini are abbreviated “N”.

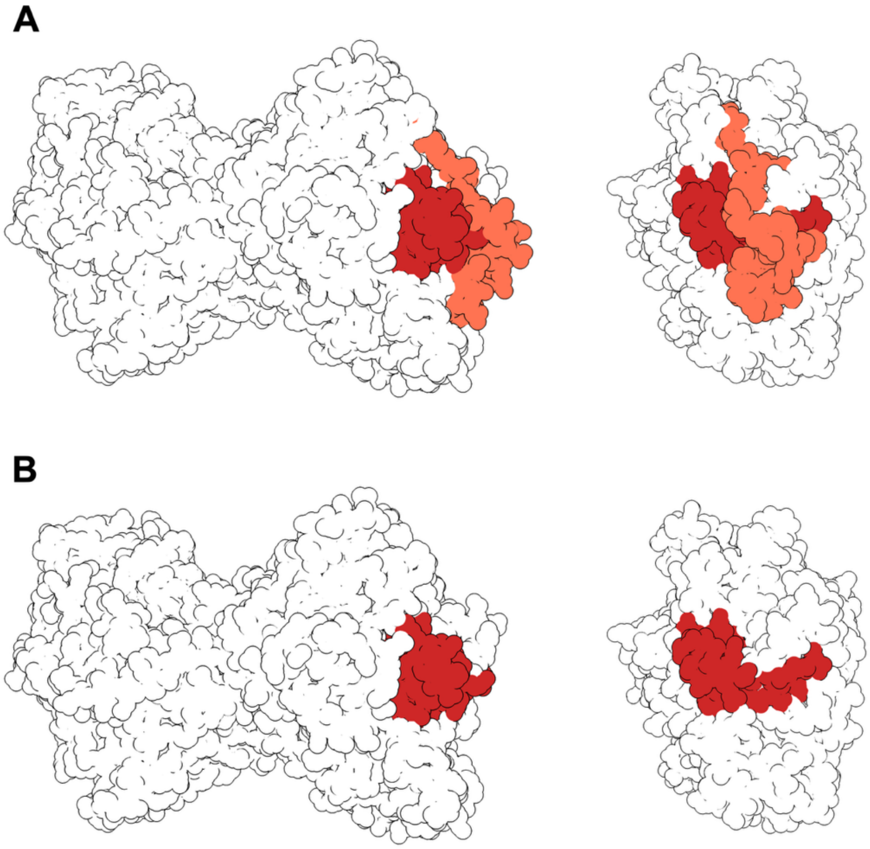


**Figure S10. The N-terminus of *Ou*IAD covers the GRD. A** When the N-terminus (colored salmon) is shown in the model of IAD, it covers the GRD (colored firebrick), akin to a seatbelt, holding the GRD inward and preventing it from moving outward for radical installation **B.** When the atoms of the N-terminus are removed from the model, the GRD is significantly more exposed and would be able to freely move out of the active site for radical installation.

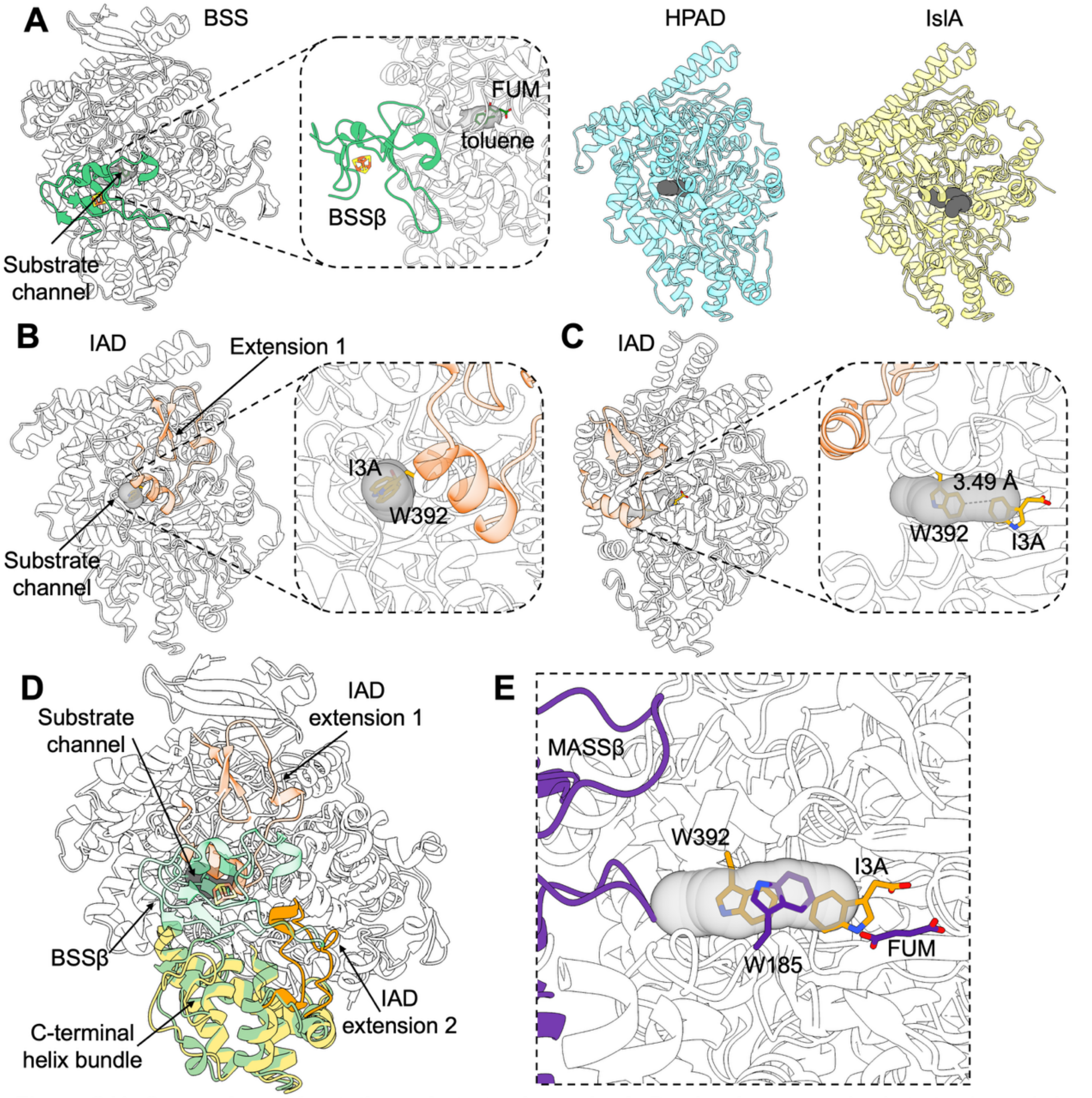


**Figure S11. Comparison of putative substrate channels. A.** Previously proposed substrate channels in BSS, HPAD, and IslA are shown. Fumarate (FUM) and toluene are shown in sticks. Channels predicted with CAVER 3.0.3. **B.** The predicted substrate channel in IAD is located near extension 1 and is blocked by W392. **C.** Side view showing W392 blocking the predicted substrate channel in IAD. **D.** Extension 2 in IAD (orange) extends from a C-terminal helix bundle (yellow) homologous to a C-terminal helix bundle in BSS (green) that was identified to move inward upon substrate and BSSβ binding. **E.** In MASS, W185 blocks the predicted substrate channel. FUM shown bound in the active site of MASS. MASSβ (rebecca purple) binds over the predicted substrate channel. Positions of IAD's W392 and substrate I3A are also shown.

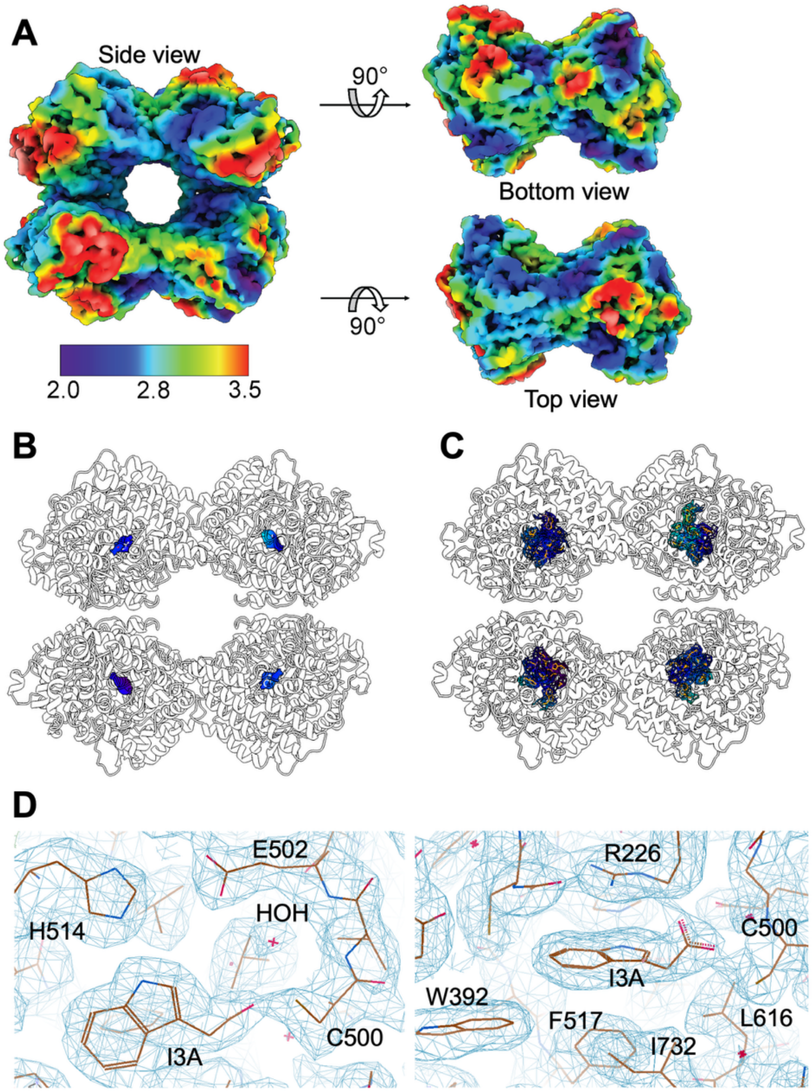


**Figure S12. Local resolution map of *Ou*IAD and resolution of active site residues. A.** Side view, top, and bottom view of the IAD tetramer. **B.** Cryo-EM map density (surface contoured to 4.5 rmsd) shows the substrate indole-3-acetate is bound in all four active sites of the tetramer and that the local resolution of all four active sites is high (between 2 and 2.8 Å resolution). **C.** Cryo-EM map density (mesh contoured to 4.5 rmsd.) shows active site residues (shown as orange sticks) visualized between 2 and 2.8 Å resolution. **D.** Views of the final IAD map and model demonstrating the clarity of resolved active site residues including those suggested to be crucial for catalysis.

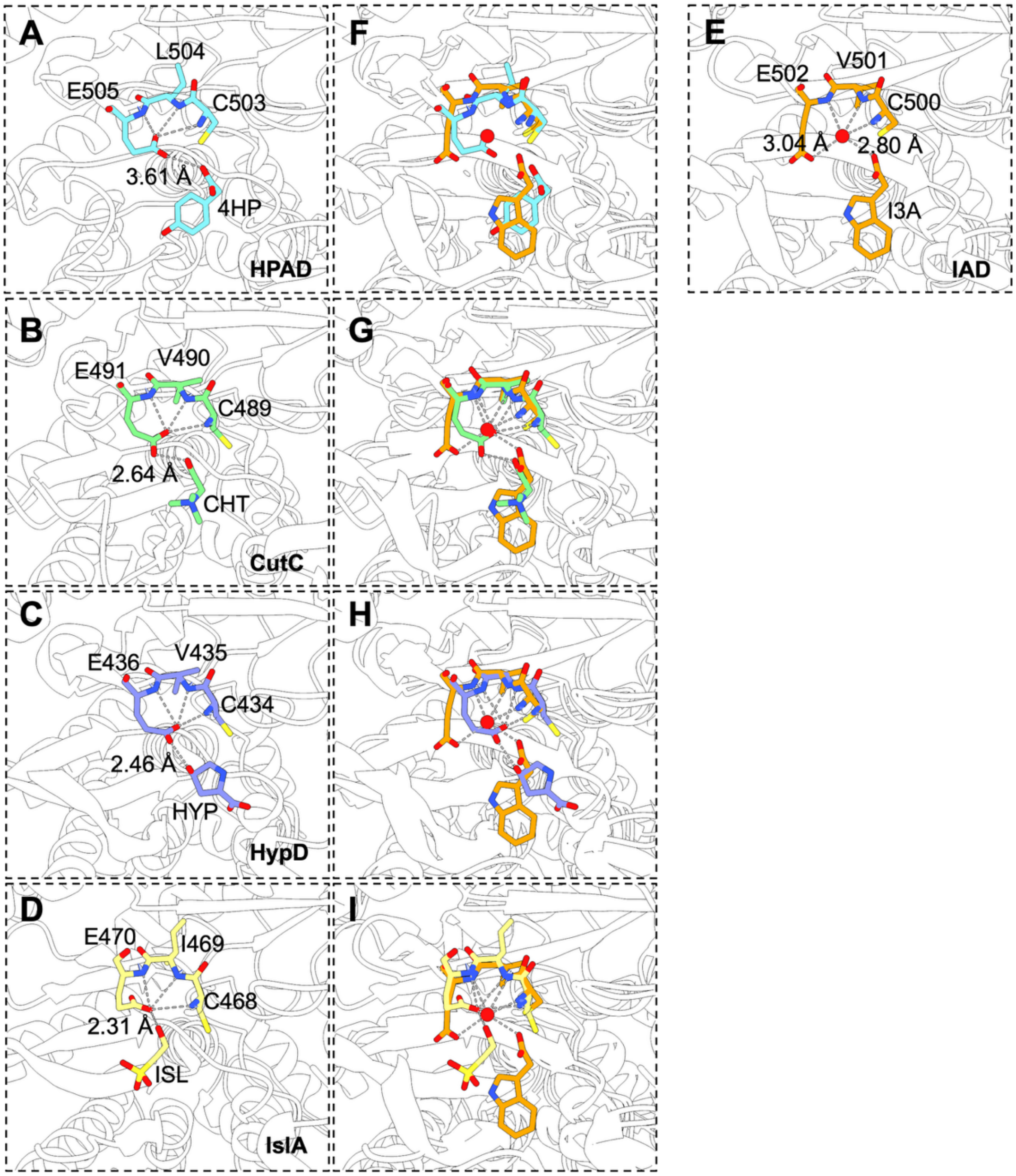


**Figure S13. CXE motif hydrogen bonding interaction with substrates of GREs. A.** CXE motif and substrate of HPAD (PDB ID: 2YAJ) **B.** CXE motif and substrate of CutC (PDB ID: 5FAU) **C.** CXE motif and substrate in HypD (PDB ID: 6VXE). **D.** CXE motif and substate in IslA (PDB ID: 7KQ3). **E.** CXE motif and substrate in IAD. Position of Glu of the CXE motif differs from that shown in A-D and a water molecule is observed in the typical position of the Glu side chain carboxylate. **F.** Overlay of IAD and HPAD (PDB ID: 2YAJ). **G.** Overlay of IAD and CutC (PDB ID: 5FAU) **H.** Overlay of IAD and HypD (PDB ID: 6VXE). **I.** Overlay of IAD and IslA (PDB ID: 7KQ3). Carbons colored by enzyme, nitrogen atoms in blue, oxygen atoms in red, and sulfur atoms in yellow.

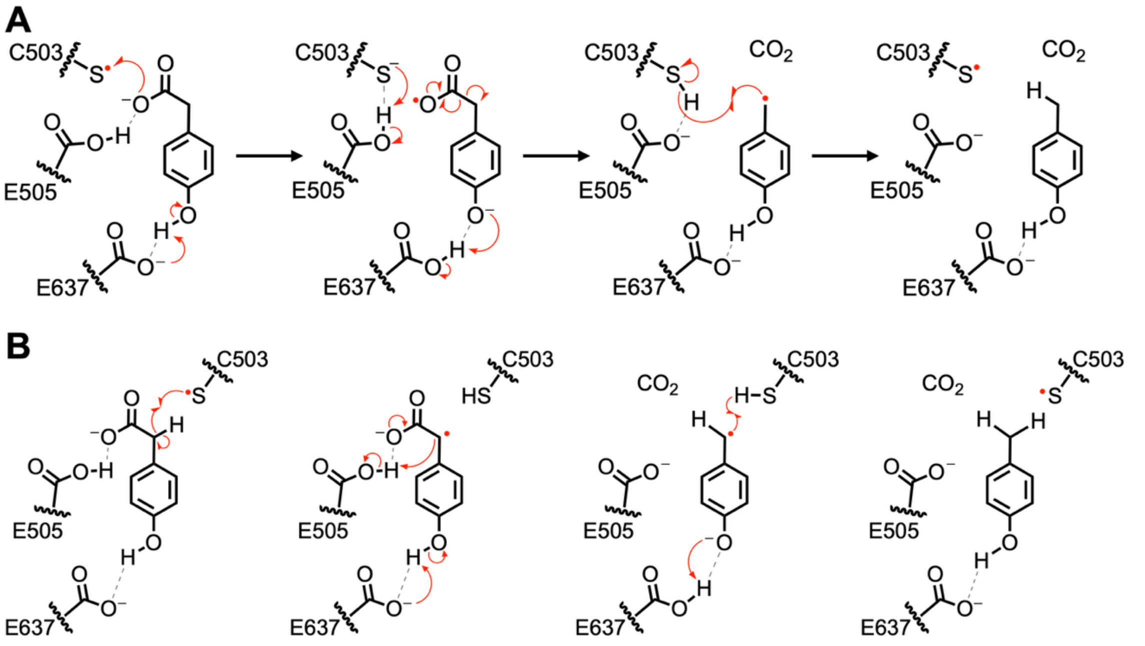


**Figure S14. Proposed mechanisms for HPAD catalysis. A.** Reaction scheme showing the proposed steps for a Kolbe-type decarboxylation mechanism catalyzed by HPAD. Briefly, the substrate carboxylate undergoes one electron oxidation by the thiyl radical coupled to deprotonation of the phenol by Glu637. The resulting thiolate is protonated by Glu505, the substrate radical readily decarboxylates, and the phenolate is reprotonated by Glu637. The product radical re-abstracts a hydrogen atom from Cys505, reforming the thiyl radical and forming product. **B.** Reaction scheme showing the steps if HPAD went by a hydrogen atom transfer mechanism.

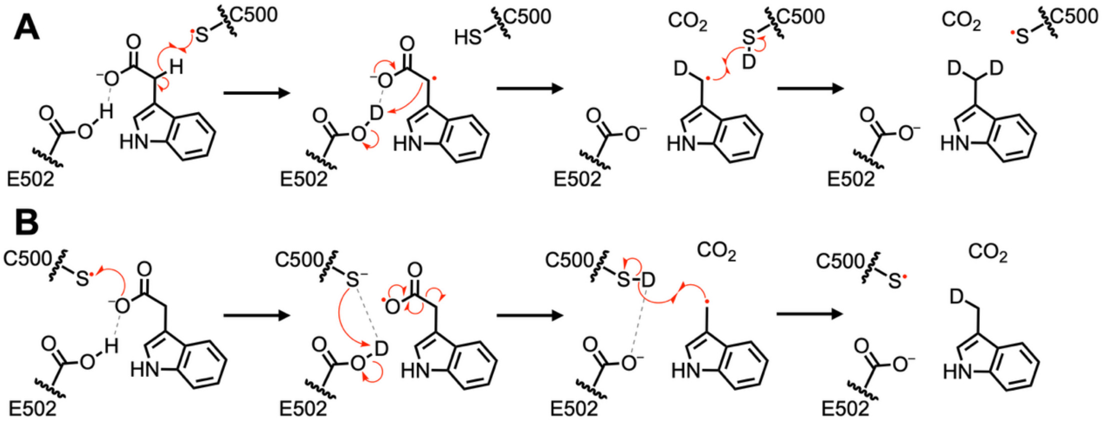


**Figure S15. Proposed mechanism of previously reported studies of *Ou*IAD deuterium incorporation into skatole product by Fu and co-workers (1). A.** Mechanism for deuterium incorporation into skatole by a HAT mechanism. Exchange of Cys500 and Glu502 with solvent allows for up to two deuteria to be incorporated into the skatole product. **B.** Mechanism for deuterium incorporation into skatole by a Kolbe-type decarboxylation. Exchange of Cys500 with solvent allows for only one deuterium to be incorporated into the skatole product.

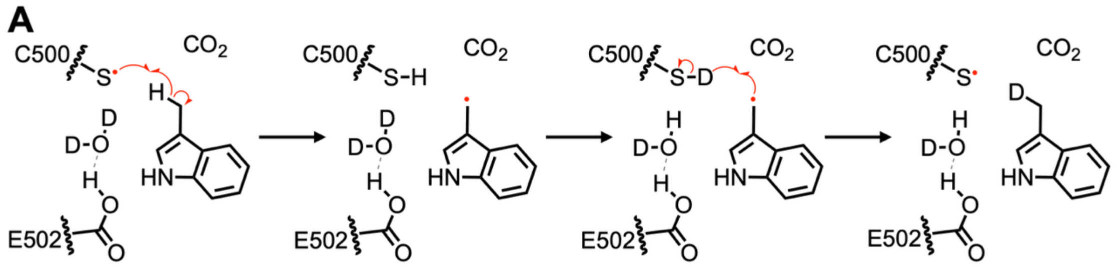


**Figure S16. Proposed mechanism of enzyme-dependent deuterium incorporation into skatole. A.** The post-turnover state of IAD is shown. The thiyl radical re-abstracts a 3'-methyl hydrogen atom from skatole. The cysteine thiol can then undergo exchange with $D_2O$, perhaps directly, or perhaps through the water molecule and Glu502. The product radical can abstract the resulting deuterium from Cys500, reforming the thiyl radical and forming deuterated product.

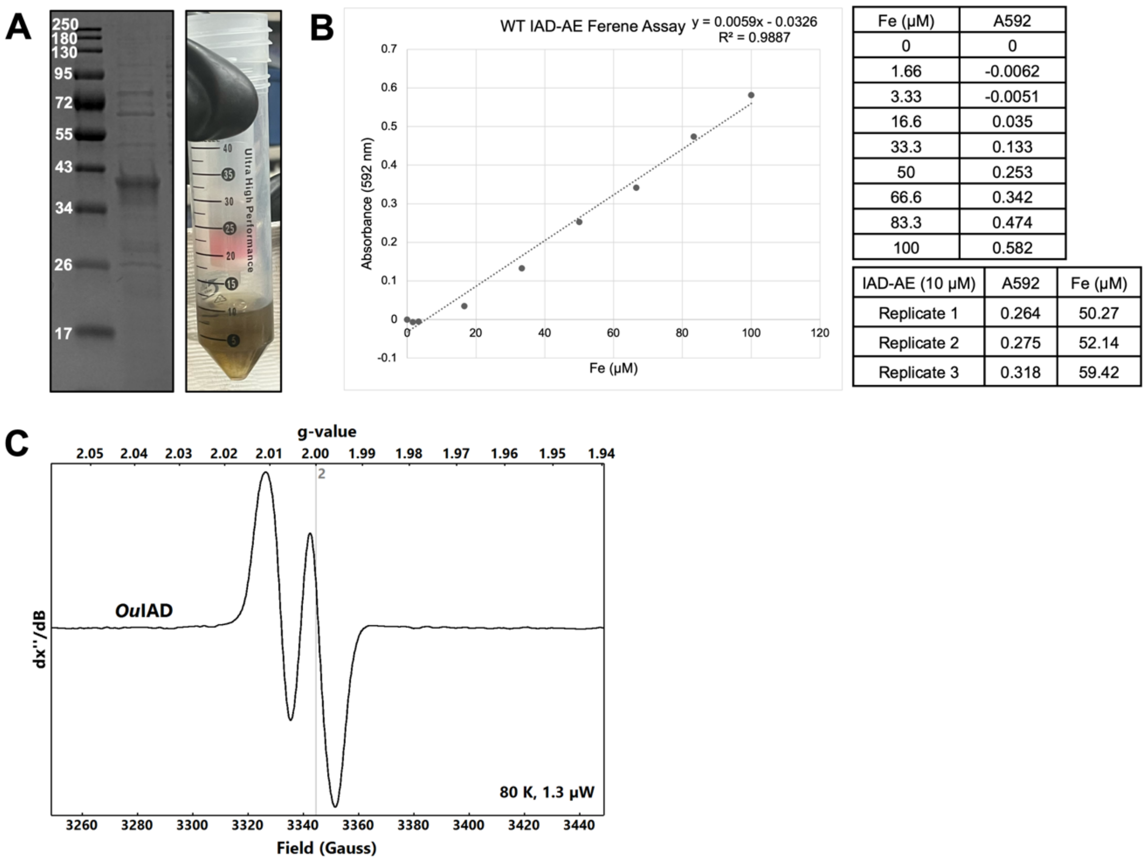


| Fe (μM) | A592 |
|---|---|
| 0 | 0 |
| 1.66 | -0.0062 |
| 3.33 | -0.0051 |
| 16.6 | 0.035 |
| 33.3 | 0.133 |
| 50 | 0.253 |
| 66.6 | 0.342 |
| 83.3 | 0.474 |
| 100 | 0.582 |

| IAD-AE (10 μM) | A592 | Fe (μM) |
|---|---|---|
| Replicate 1 | 0.264 | 50.27 |
| Replicate 2 | 0.275 | 52.14 |
| Replicate 3 | 0.318 | 59.42 |

**Figure S17. Purification of *Ou*IAD-AE, activation of *Ou*IAD, and EPR spectrum of a *Ou*IAD glycyl radical. A.** SDS-PAGE gel showing purified 6xHis-*Ou*IAD-AE (37.18 kDa) **B.** Fe standard curve and Fe analysis of purified *Ou*IAD-AE protein shows 5.39 ± 0.48 Fe per monomer. **C**. Characteristic doublet signal at g=2.0 of the glycyl radical, demonstrating *Ou*IAD-AE can install 0.52 radicals per *Ou*IAD monomer.

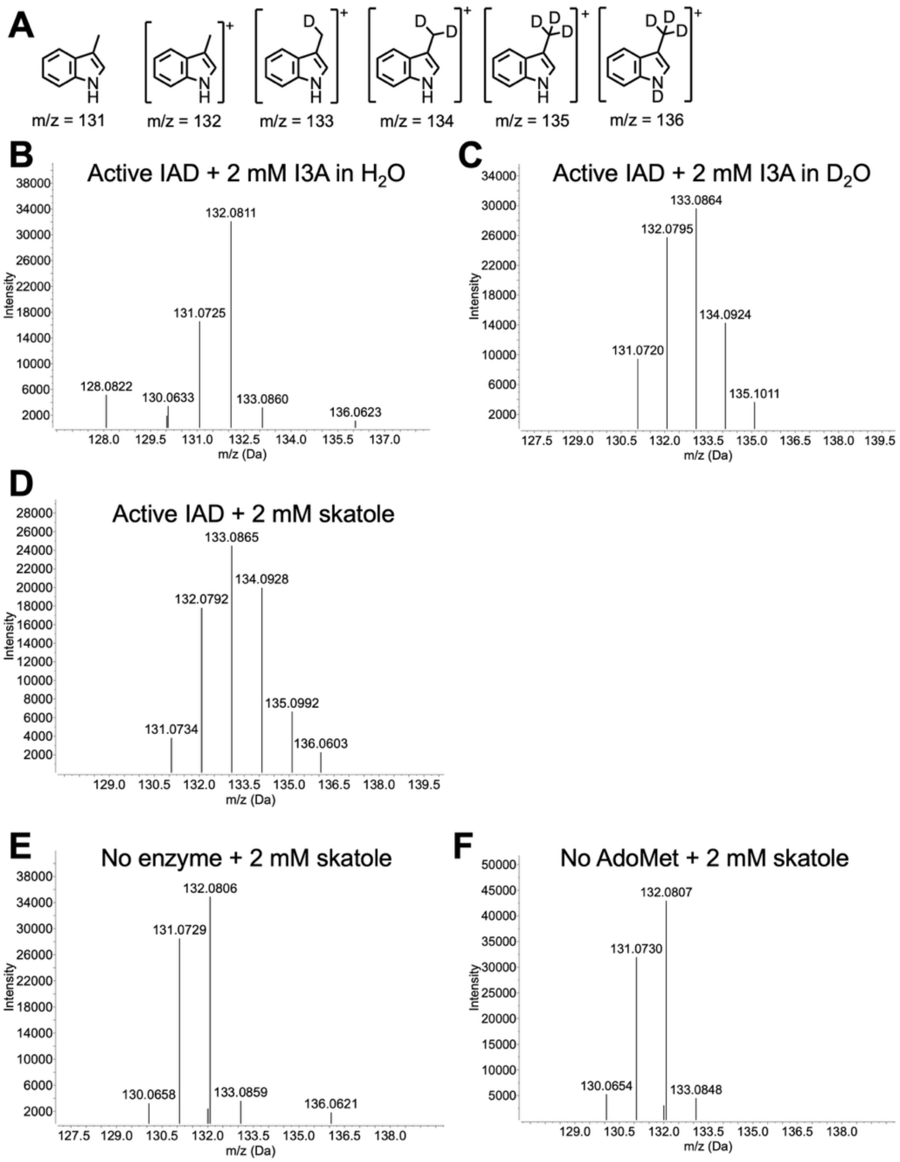


**Figure S18. *Ou*IAD *in vitro* activity on its native product skatole. A.** Expected masses in positive ion mode of deuterated skatole. **B.** Activated *Ou*IAD can turnover the native substrate indoleacetate in $H_2O$ to produce skatole. Some amount of indole N–H to N–D exchange may also occur. **C.** When *Ou*IAD is activated in $D_2O$ buffer and provided with indoleacetate, the products skatole, $D_1$- $D_2$- and $D_3$-skatole are observed consistent with previous results by Fu *et al*. (1). **D.** The thiyl radical of *Ou*IAD can act on skatole, exchanging 3′-methyl hydrogens with deuteria to produce $D_1$-skatole and $D_2$-skatole, as observed by LC/MS. **E.** No enzyme control shows that no deuterium incorporation into skatole is observed when enzyme is omitted from assays. **F.** No AdoMet control shows no deuterium incorporation into skatole is observed when AdoMet is omitted from assays.

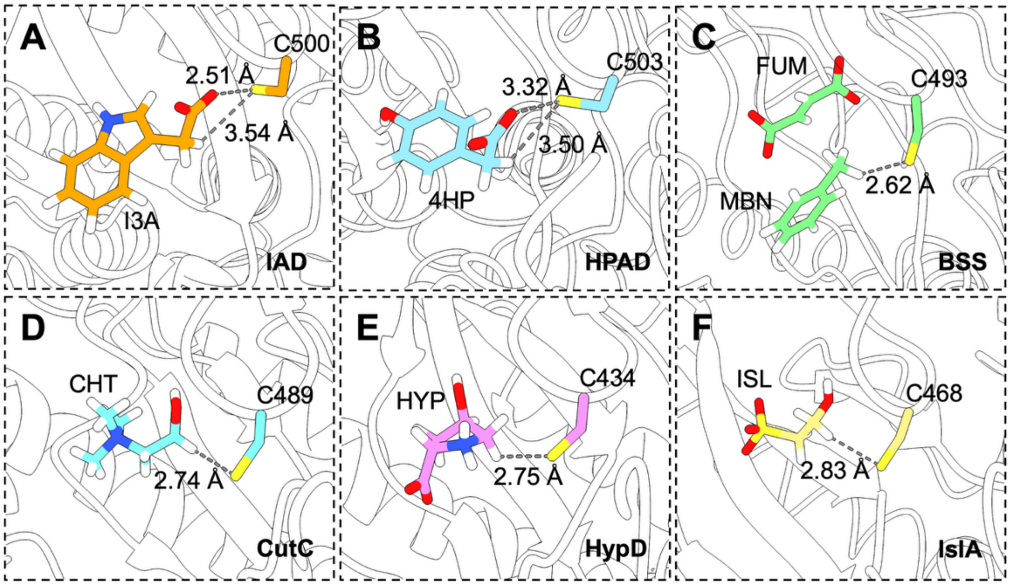


**Figure S19. The catalytic cysteine in GREs is oriented toward the site of hydrogen atom abstraction. A.** In IAD, the catalytic Cys500 is not oriented toward a hydrogen atom of the substrate indoleacetate (I3A) for abstraction. **B.** Similarly, in HPAD, Cys503 is not oriented toward a hydrogen atom of the substrate hydroxyphenylacetate (4HP) for abstraction. **C.** In BSS, Cys493 is oriented toward a hydrogen atom of the substrate toluene (MBN) for abstraction. **D.** In CutC, Cys489 is oriented toward a hydrogen atom of the substrate choline (CHT) for abstraction. **E.** In HypD, Cys434 is oriented toward a hydrogen atom of the substrate hydroxyproline (HYP) for abstraction. **F.** In IslA, Cys468 is oriented toward a hydrogen atom of the substrate isethionate (ISL) for abstraction. Hydrogen atoms were added in Coot (8).

**Table S1.** Cryo-EM data collection, reconstruction, and model refinement of I3A-bound *Ou*IAD

| | |
|---|---|
| **Data collection** | |
| Microscope | FEI Titan Krios |
| Camera | G3i/BioQuantum K3 |
| Acceleration voltage (kV) | 300 |
| Spherical aberration (mm) | 2.7 |
| Magnification (x) | 105,000 |
| Pixel size (Å) | 0.822 |
| Defocus range (μm) | -0.2 to -2.0 |
| Number of frames | 50 |
| Exposure time (s) | 1.9 |
| Total exposure ($e^-$/Å$^2$) | 52.07 |
| Shots per hole | 7 |
| Total micrographs collected | 8927 |
| Total particles | 2743992 |
| Automation software | EPU |
| **Reconstruction** | |
| Software | RELION-4.0 |
| Final number of particles | 330270 |
| Symmetry imposed | none |
| Unmasked resolution at 0.143 FSC (Å) | 2.74 |
| Masked resolution at 0.143 FSC (Å) | 2.29 |
| Local resolution range | 2.33-4.36 |
| **Model refinement** | |
| Refinement software | Phenix |
| Refined model resolution at 0.143 FSC (Å) | 2.1 |
| $CC_{volume}$/$CC_{mask}$/$CC_{box}$ | 0.89/0.90/0.89 |
| Mean CC for ligands | 0.71 |
| Model composition | |
| Non-hydrogen atoms | 28486 |
| Residues | 3500 |
| Indole-3-acetate (IAC) | 4 |
| Water | 686 |
| Rms deviations | |
| Bond lengths (Å) | 0.003 |
| Bond angles (deg) | 0.744 |
| MolProbity score (11) | 1.74 |
| Clash score | 9.38 |
| Rotamers outliers (%) | 1.39 |
| Ramachandran (%) | |
| Favored | 97.31 |
| Allowed | 2.46 |
| Outliers | 0.23 |
| CaBLAM outliers (%) | 1.38 |
| EMRinger score (12) | 4.68 |
| Average Q-score | |
| Protein | 0.814 |
| IAC | 0.849 |
| Waters | 0.894 |
| B-factors (min/max/mean) | |
| Protein | 4.20/96.94/27.26 |
| IAC | 9.89/19.58/13.28 |
| Waters | 4.20/54.61/21.68 |
